# Dysregulation of extracellular vesicle protein cargo in female ME/CFS cases and sedentary controls in response to maximal exercise

**DOI:** 10.1101/2023.08.28.555033

**Authors:** Ludovic Giloteaux, Katherine A. Glass, Arnaud Germain, Sheng Zhang, Maureen R. Hanson

**Author notes:** These authors contributed equally and are listed alphabetically (Ludovic Giloteaux, Katherine A. Glass). Corresponding author: Maureen R. Hanson, Department of Molecular Biology and Genetics, Cornell University, 323 Biotechnology Building, 526 Campus Road, Ithaca, New York 14853, USA.

## Abstract

In healthy individuals, physical exercise improves cardiovascular health and muscle strength, alleviates fatigue, and reduces risk of chronic diseases. Although exercise is suggested as a lifestyle intervention to manage various chronic illnesses, it negatively affects people with myalgic encephalomyelitis/chronic fatigue syndrome (ME/CFS), who suffer from exercise intolerance. We hypothesized that altered extracellular vesicle (EV) signaling in ME/CFS patients after an exercise challenge may contribute to their prolonged and exacerbated negative response to exertion (post-exertional malaise). EVs were isolated by size exclusion chromatography from the plasma of 18 female ME/CFS patients and 17 age- and BMI-matched female sedentary controls at three time points: before, 15 minutes, and 24 hours after a maximal cardiopulmonary exercise test. EVs were characterized using nanoparticle tracking analysis and their protein cargo was quantified using Tandem Mass Tag-based (TMT) proteomics. The results show that exercise affects the EV proteome in ME/CFS patients differently than in healthy individuals and that changes in EV proteins after exercise are strongly correlated with symptom severity in ME/CFS. Differentially abundant proteins in ME/CFS patients vs. controls were involved in many pathways and systems, including coagulation processes, muscle contraction (both smooth and skeletal muscle), cytoskeletal proteins, the immune system, and brain signaling.

## INTRODUCTION

Myalgic Encephalomyelitis/Chronic Fatigue Syndrome (ME/CFS) is a complex and highly disabling chronic illness that affects millions of people worldwide. It can affect individuals of any age, gender, or socioeconomic background and is characterized by persistent fatigue that is not relieved by rest, along with a range of other symptoms including muscle and joint pain, cognitive difficulties, sleep disturbances, and headaches (IOM, 2015). While the exact cause of ME/CFS remains unknown, it is believed to involve a complex interplay of genetic, environmental, and immunological factors.

One of the hallmark features of ME/CFS is post-exertional malaise (PEM), which is an exacerbation of symptoms that occurs after even minimal exertion due to basic activities of daily living, physical activity, cognitive work, or engaging in social interactions. The intensity and duration of PEM is usually unpredictable, making it challenging for individuals with ME/CFS to plan activities or engage in regular daily tasks (IOM, 2015; Stussman et al., 2020). Research suggests that PEM may involve dysregulation of the immune system, impaired energy metabolism, and abnormal responses to exertion at the cellular level (Germain et al., 2022; Glass et al., 2023; Van Booven et al., 2023).

In healthy individuals, physical exercise improves cardiovascular health and muscle strength, alleviates fatigue, anxiety and depression, and reduces risk of chronic diseases (Nystoriak & Bhatnagar, 2018; Singh et al., 2023). Although exercise is suggested as a lifestyle intervention to manage various chronic illnesses (Li et al., 2023; Puetz & Herring, 2023; Reina-Gutiérrez et al., 2023; Ross et al., 2023), it negatively affects people with ME/CFS who suffer from exercise intolerance. This intolerance has been extensively documented using a two-day cardiopulmonary exercise test (CPET), with the aim to understand and characterize the specific physiological responses and limitations that individuals with ME/CFS experience during and after exercise (Keller et al., 2014; Moore et al., 2023; Stevens et al., 2018).

The health benefits of exercise have been attributed in part to bioactive molecules (metabolites, RNA species, and proteins) released into circulation during exercise (Contrepois et al., 2020). These molecules may play a role in the response to exercise by triggering the required adaptations and/or regulating homeostasis and substrate metabolism (Murphy et al., 2020). Such systemic mediators need to be targeted to exert their biological effects, and circulating extracellular vesicles (EVs) can facilitate cell-to-cell communication, as well as organ cross-talk both locally and at distant sites through receptor-ligand interactions and subsequent release of their cargo (Gurung et al., 2021). EVs are spherical entities protected by a lipid bilayer that range from 30 – 1,000 nm in diameter and are detected in most body fluids including blood, urine, and saliva. EVs are secreted by skeletal muscle, endothelial, cardiac, hepatic, and adipose tissues as well as immune cells, especially platelets (Doyle & Wang, 2019; Haghighitalab et al., 2023; Kehrloesser et al., 2023). The cargo of EVs released after exercise reflects the cellular response to exercise-induced stress including alterations in metabolism, inflammation, and tissue repair (Whitham et al., 2018). Athletic males exhibit an increase in the concentration of circulating EVs during and immediately after an acute bout of aerobic exercise and analysis of the EV protein cargo showed changes in numerous proteins associated with various signaling pathways including angiogenesis, immune signaling, and glycolysis (Frühbeis et al., 2015; Whitham et al., 2018). Regrettably, there is limited research regarding the effects of physical activity on EVs in disease populations, healthy females, or sedentary individuals.

To date, few studies on EVs in ME/CFS have been published (Almenar-Pérez et al., 2020; Bonilla et al., 2022; Castro-Marrero et al., 2018; Eguchi et al., 2020; Giloteaux et al., 2023; Giloteaux et al., 2020; González-Cebrián et al., 2022). In 2018, Castro-Marrero and colleagues were the first to analyze EVs isolated from ME/CFS patients and they found a higher EV concentration and smaller sized EVs in ten ME/CFS patients in comparison to five healthy controls (Castro-Marrero et al., 2018). Almenar-Pérez et al. replicated these results and found dysregulated neuroimmune pathways via analysis of EV miRNA content in a study with 15 severely affected ME/CFS patients compared to 15 healthy subjects (Almenar-Pérez et al., 2020). Our group published two studies measuring the cytokine content of EVs isolated from plasma of 35 and 49 ME/CFS patients and matched healthy controls, respectively (Giloteaux et al., 2023; Giloteaux et al., 2020). Although we did not replicate EV size differences between groups, all four studies found significantly higher concentration of EVs in the ME/CFS group. The cytokine analysis revealed few significant differences compared to controls but significant correlations with patient symptom severity. Eguchi and colleagues also replicated the finding of increased concentration of circulating EVs in a larger cohort of 99 ME/CFS patients compared to 53 controls (Eguchi et al., 2020). However, one study in which EVs were directly quantified without prior isolation showed no difference in EV concentration between 20 ME/CFS patients and 20 controls (Bonilla et al., 2022). Further proteomic analysis of EV cargo in addition to cytokines could help identify a signature biomarker for ME/CFS. Quantitative untargeted proteomics of isolated EVs from three ME/CFS patients and three controls from the larger Eguchi et al. cohort revealed a protein cargo specific to the ME/CFS group including actin network proteins and 14–3-3 family proteins (Eguchi et al., 2020), but these results need to be validated in a larger study.

We hypothesize that altered EV signaling in ME/CFS patients after an exercise challenge may contribute to the prolonged and exacerbated negative response to exertion in ME/CFS (PEM). It is possible that EVs are carrying aberrant signaling molecules, that signals that should be transmitted in EVs post-exercise are missing in ME/CFS patients, and/or that the EV response to exercise is temporally dysregulated.

In the present study, we isolated EVs by size exclusion chromatography from the plasma of 18 females diagnosed with ME/CFS and 17 age- and Body Mass Index (BMI)-matched female sedentary controls at three time points: before, 15 minutes, and 24 hours after a maximal exercise challenge. We then characterized the EVs using nanoparticle tracking analysis (NTA) and measured the protein cargo using Tandem Mass Tag-based (TMT) quantitative proteomics. We demonstrate that exercise affects the EV proteome in ME/CFS patients differently than in healthy individuals and that changes in EV proteins after exercise are strongly correlated with symptom severity in ME/CFS.

## MATERIALS AND METHODS

### Population characteristics

A cohort of 18 females diagnosed with ME/CFS and 17 female age- and BMI-matched sedentary controls (Table 1) was selected for this study (a subset of the larger cohort, ClinicalTrials.gov Identifier: NCT04026425). All cases met the 2003 Canadian Consensus Criteria for ME/CFS (Carruthers et al., 2003). None of the subjects in our cohort were pregnant or breastfeeding, diabetic, smoked cigarettes, consumed excessive amounts of alcohol, had an orthopedic limitation preventing them from performing the cardiopulmonary exercise test (CPET), or had any of the following diagnoses: schizophrenia, major depressive disorder, bipolar disorder, or an anxiety disorder. Autoimmune compromised controls were also excluded.

**Table 1:** Study population characteristics.

|  |  | ME/CFS | Controls | p-value |
| --- | --- | --- | --- | --- |
| n |  | 18 | 17 | NA |
| Age (years) |  | 44.4 ± 7.9 | 44.2 ± 13.4 | 0.87 |
| BMI |  | 27.8 ± 5.4 | 29.9 ± 5.8 | 0.26 |
| Type of onset | Sudden | 83% | NA | NA |
|  | Gradual | 17% | NA | NA |
| Bell Disability Scale | Median (IQR) | 30 (17.5) | 100 (10) | p < 0.0001 |
| SF-36 PCS | Physical Component Summary | 25.2 ± 6.7 | 56.2 ± 5.0 | p < 0.0001 |
| SF-36 MCS | Mental Component Summary | 46.0 ± 8.1 | 54.5 ± 6.9 | p = 0.0001 |
| MFI total | Multi-dimensional fatigue inventory | 84.4 ± 7.7 | NA | NA |

Shown are the mean and standard deviation for demographic parameters in ME/CFS patients and control subjects, except for the Bell's disability scale. For the Bell disability scale and SF-36, a higher number corresponds to better health. For the multi-dimensional fatigue inventory (MFI) total score, a higher number corresponds to increased fatigue. The p-values are from a Mann Whitney U test. NA: not applicable.

### Exercise testing and blood sample collection

All blood samples were collected prior to March 2020, allowing us to distinguish ME/CFS cases from people with post-acute sequelae of COVID-19 (Long COVID). Blood was collected at baseline (0h), immediately prior to a CPET. All subjects performed a maximal CPET on a stationary bicycle by cycling with an increase in workload of 15 watts per minute of exercise until volitional exhaustion, with a respiratory exchange ratio greater than 1.1 (the threshold for maximum effort). A second blood sample was also collected approximately 15 minutes post-CPET (15min) and a third one the following day (24h). Peripheral blood was drawn in sodium citrate BD Vacutainer^TM^ Cell Preparation Tubes and centrifuged to pellet red blood cells.

Resulting plasma samples were stored at −80°C prior to processing. Baseline health status was assessed using the Short Form 36 Health Survey v2® (SF-36) (Ware, 1993), the Bell Disability Scale, and the Multidimensional Fatigue Inventory (MFI) (Smets et al., 1995). A modified version of the chronic fatigue syndrome specific symptom severity (SSS) score survey (Baraniuk et al., 2013) was used to assess symptom severity at different time points, including on the days of blood collection. Written informed consent was obtained from all participants and all protocols were approved by the Ithaca College Institutional Review Board (protocol 1017-12Dx2), and the Weill Cornell Medical College Institutional Review Board (protocol 1708018518). The research adhered to the tenets of the Declaration of Helsinki.

### Extracellular vesicles purification and characterization

Extracellular vesicles (EVs) were isolated from 500 µL of plasma by size exclusion chromatography using Izon qEVoriginal/70 nm columns (Izon Science) following the manufacturer’s instructions. Halt™ Protease Inhibitor Cocktail (1X) (Thermo Scientific™) was added to samples prior to submission for proteomic analysis (see below). Concentration and size distribution of isolated EVs were assayed in all samples using a NanoSight NS300 instrument for nanoparticle tracking analysis (NTA) (Malvern, Worcestershire, UK). Samples were thawed and diluted to 1:2000 in PBS 1X and 1 mL was injected through the laser chamber. Three recordings of 60-second digital videos of each sample were acquired and analyzed by the NanoSight NTA 2.3 software to determine the size and the concentration of EVs. The non-parametric Wilcoxon rank sum test was used to test the significance of differences (p < 0.05) between cases and controls for the size and EV concentration. The Wilcoxon signed rank test was used to statistically compare the size and concentration at different time points within groups (p < 0.05). EV suspensions were visualized by Transmission Electron Microscopy (TEM). Undiluted samples were applied to copper 300-mesh Formvar coated carbon stabilized TEM grids (Electron Microscopy Sciences) and were allowed to adsorb to the grid for 5 minutes. Grids were then washed twice by floating on distilled water for 30 seconds. Negative staining was performed by placing the grids on a drop of 2% Aqueous Uranyl Acetate for 10 minutes. The grids were blot dried with Whatman^TM^ paper and imaged with a JEOL JEM 1230 Transmission Electron microscope (JEOL USA, Inc.).

### Proteomics experimental procedures

Detailed information can be found in supplementary materials (Supplementary File S1). A Tandem Mass Tag (TMT) 10-plex shotgun proteomics analysis was used in this study (Supplementary Figure 1). Specifically, nine protein-extracted EV samples from three different subjects at 0h (baseline), 15 minutes, and 24 hours post-exercise were included in each of two TMT 10-plex sets for the ME/CFS group and the control group, respectively. The remaining one channel in each TMT 10-plex set was used for a mixture (pooled reference) containing equal amounts of proteins from each of the 18 samples to bridge the results between the control and ME/CFS groups. A total of 6 TMT experiments were conducted generating 12 datasets. Fifteen µg of protein in each sample was reduced, alkylated and digested with 25 µL trypsin at 60 ng/µL in 50 mM Triethylamonium bicarbonate (TEAB) buffer pH 8.5 in a S-Trap Micro Spin column (Protifi, Huntington, NY). Further labeling of the tryptic digest samples was performed according to Thermo Scientific’s TMT Mass Tagging Kits and Reagents protocol. Ten labeled samples in each of TMT 10-plex sets were pooled together and 80 µg of labeled tryptic peptides were then fractionated using a Pierce™ High pH Reversed-Phase Peptide Fractionation Kit (Thermo-Fisher Scientific, San Jose, CA) with a 9-step isocratic elution. The final three fractions were pooled and subjected to nanoLC-MS/MS analysis using an UltiMate3000 RSLAnano coupled with Orbitrap Eclipse mass spectrometer (Thermo-Fisher Scientific, San Jose, CA) operated under the Real Time Research (RTS) SPS MS3 method (Fu et al., 2021). All data was acquired and processed with the Thermo Scientific™ Xcalibur™ software 4.3. Acquired raw MS files were processed and searched using the Sequest HT search engine within the Proteome Discoverer 2.3 software (PD 2.3, Thermo).

The MS/MS spectra were searched against the *Homo Sapiens* NCBI database downloaded on 3/14/2019 (81,268 sequences). Oxidation of Met (M), deamidation of Asn (N) and Gln (Q), protein N-terminal acetylation, M-loss and M-loss+acetylation were set as variable modifications; carbamidomethyl Cys (C) and TMT 10-plex (m/z 229.1629) on peptide N-terminal and Lys (K) residue were specified as a static modification. The TMT quantification workflow within the PD software was used for identity of peptides and to calculate the reporter ion abundances in MS3 spectra corrected for isotopic impurities. Identified peptides were filtered for maximum 1% FDR using the Percolator algorithm in PD 2.3 with peptide confidence set to high and peptide mass accuracy ≤5 ppm. Both unique and razor peptides were used for quantitation. Signal-to-noise (S/N) values were used to represent the reporter ion abundance with a co-isolation threshold of 50% and an average reporter S/N threshold of 10. The S/N values of peptides, which were summed from the S/N values of the peptide-spectrum matches (PSMs), were summed to represent the abundance of the proteins. For relative ratio between the two groups, normalization on total peptide amount for each sample was applied. Comparison of ratios of each sample vs. the pooled reference allows protein abundance across two different TMT sets to be assessed (Unwin et al., 2010).

### Data processing and statistical analysis

A total of 862 proteins were detected in the six TMT experiments. Lipoproteins and keratins were removed prior analysis as known contaminants in EV proteomics studies (Karimi et al., 2018; Théry et al., 2018). 301 proteins remained for analysis after filtering out proteins that were not detected in at least 72 out of 108 samples and 4 out of 6 TMT experiments (two thirds, see Supplementary File S2). For these 301 proteins, missing values were then imputed with random forest (RF, *missForest* R package, default parameters (Stekhoven & Bühlmann, 2012). All proteins and the following categorical variables with no missing data were included in the RF imputation: experimental group (ME/CFS vs. Control), time point (0h, 15min and 24h) and TMT experiment (sets 1-6).

Original and imputed data for 301 EV proteins were analyzed at all three time points. Due to the experimental setup, ME/CFS and control samples could only be compared within each TMT experiment. Therefore, we used a bootstrapping approach to statistically compare ME/CFS and control samples at each time point. For each protein, the following algorithm was used to generate 10,000 bootstrapped datasets: 1) select 18 values with replacement, 3 per TMT experiment, for ME/CFS and control groups, 2) calculate 18 ME/CFS vs. control fold changes from the selected values (all fold changes are matched within experiments) and 3) calculate the median of the fold changes, which is less influenced by outliers than the mean. To compare the ME/CFS to the control group, 95% confidence intervals were calculated for the 10,000 bootstrapped medians. The null hypothesis is that the ratio of the two groups is 1 (no difference). Confidence intervals were adjusted to account for false discovery using the Benjamini and Yekutieli procedure (q < 0.1) (Benjamini & Yekutieli, 2005). Protein levels are considered significantly different between groups if the adjusted confidence interval does not include 1. Those proteins are differentially abundant proteins (DAPs). Therefore, if the confidence interval is below 1, that DAP is significantly lower in ME/CFS compared to controls and if the confidence interval is above 1, then the DAP is significantly higher in the ME/CFS group.

To compare DAPs over time within the ME/CFS and control groups, we calculated the within-subject fold changes for 15min vs 0h, 24h vs 0h, and 24h vs 15min. This analysis did not require a bootstrapping approach because all three samples for every subject were measured within the same TMT experiment. For each protein, the mean fold changes were compared to 1 using a t-test. Again, the null hypothesis was that the mean was equal to 1 (no difference). The p-values were adjusted using the Benjamini and Hochberg false discovery rate (BH FDR) correction procedure (q < 0.15). For significantly different proteins, if the mean fold change is above 1, then that protein increased over the two time points compared and if the mean fold change is below 1, then the level of that protein decreased over time.

To assess whether the protein levels was changing differently over time in the ME/CFS vs. control groups, we used a similar bootstrapping approach to that described above. For this analysis, we randomly selected 18 within-subject fold changes with replacement, 3 per TMT experiment, for both the ME/CFS and control groups. Steps 2 and 3 of the algorithms outlined above were the same, and we followed the same statistical procedure. Proteins that are significantly different in this analysis again had adjusted confidence intervals that did not include 1. In this case, the median fold change is simply telling us the difference between the change over time in ME/CFS vs. controls, but not the direction of that change within each group.

### Pathway enrichment analyses

Significant DAPs were subjected to functional annotation of Reactome pathway enrichment analysis using the ShinyGO (v0.76) software (Ge et al., 2020). Reactome terms and pathways with adjusted p-value FDR < 0.05 were considered as significantly enriched. FDR is calculated based on nominal p-value from the hypergeometric test.

Canonical pathway analysis was performed with QIAGEN IPA software (QIAGEN Inc.). For each time point, the input included all 301 proteins and the median fold change of patients vs. controls generated from the 10,000 bootstrapped datasets. 295 proteins with valid IDs in IPA were preserved for further steps. Because all proteins from the dataset were supplied for each time point, enrichment significance (p-values) was identical for all time points. The Fisher’s Exact Test was used to calculate a statistical significance (p-value) of overlap of the dataset molecules with various sets of molecules that represent annotations such as canonical pathways. Subsequently, p-values were adjusted using the BH FDR procedure. The overall activation/inhibition states of canonical pathways are predicted based on a z-score algorithm using the median fold changes for the proteins in each pathway. The primary purpose of the activation z-score is to infer the activation states (“increased” or “decreased”) of implicated biological functions in the two groups being compared (i.e. ME/CFS patients vs controls). The sign of the calculated z-score reflects the overall predicted activation state of the biological function (<0: inhibited, >0: activated). An absolute z-score of ≥ 2 is considered significant (Krämer et al., 2014). After first sorting by BH FDR corrected p-values, the top 30 canonical pathways were retained and hierarchically clustered by z-score where six pathways without a z-score were eliminated.

### Tissue and cell type enrichment analysis

Tissue and cell type enrichment analysis was performed using the DAPs within each group 15 minutes post-exercise (q < 0.15). The 187 DAPs in controls and the 63 DAPs that were significantly increased in ME/CFS patients were compared with the following databases in the tissue enrichment subsection of Enrichr: Human Gene Atlas, ARCHS4 Tissues, and HuBMAP ASCT+ B augmented with RNAseq coexpression (Chen et al., 2013; Xie et al., 2021). Fisher’s exact test was used to test whether the proteins that changed after exercise were significantly overlapping with the annotated gene/protein sets, and p-values were adjusted with the BH FDR correction procedure (significance threshold q < 0.15). We identified the tissue and cell type terms that were uniquely enriched in either the control group or the ME/CFS group, and the terms that were significantly enriched in both groups. Since this analysis allows for overlap (meaning each protein can contribute to the enrichment of several terms), Sankey networks were constructed to show all proteins that contributed to the tissue terms unique to the ME/CFS or control groups (*networkD3 package*, https://cran.r-project.org/web/packages/networkD3/index.html). The tissue and cell type terms were grouped in physiologic categories for ease of visualization in the Sankey networks. See Supplementary Tables S8, S9 and S10) for the original tissue and cell type enriched terms, the databases they belong to, and the category assignments.

### Correlation with clinical parameters

To investigate whether changes in protein abundance after exercise are associated with ME/CFS symptoms, severity metrics, and demographic parameters, Spearman correlation coefficients for the within-subject fold changes over time for each protein vs. a variety of clinical parameters were calculated. Significance was assessed by generating bootstrapped 95% confidence intervals for each correlation coefficient and getting associated p-values (*bootcorci* R package) with the null hypothesis that the correlation coefficient equals 0. Similar to the previous bootstrapping analysis, the correlation is significant if the 95% confidence interval does not include 0. For each clinical parameter, p-values were adjusted using BH FDR correction (q < 0.1). All three time point ratios were evaluated (15min/0h, 24h/0h, and 24h/15min). We define significant and strong correlations as an absolute value of Spearman R > 0.7 and q < 0.1.

Demographic parameters included age, BMI, and ME/CFS duration (ME/CFS group only). Survey data that reflects physical function and disease severity were also evaluated including the Bell Disability Scale score, the SF-36 Physical Component Summary (PCS), and the percentage of waking time spent in a reclined position. The MFI-20 total score was also used as another metric of general fatigue in the ME/CFS subjects only (range 20 – 100, higher scores correspond to higher levels of fatigue).

Furthermore, we looked at symptom severity using the SSS scores. Each subject in both the ME/CFS and control groups was asked to rate their symptoms on a scale of 0-10 (0 = not present and 10 = very severe) at three different times: 1) on average over the past month (Initial), 2) on the morning of the CPET (0h) and 3) 24 hours after the CPET (24h). The ΔSSS are calculated by the score for the symptom at 24h minus the score at 0h. Thus, ΔSSS below 0 indicates an improvement in that symptom following exercise and ΔSSS above 0 indicates that the symptom got worse 24h post-exercise. The following SSS were included in our analysis: fatigue, impaired memory or concentration, recurrent sore throat, lymph node tenderness, muscle tenderness or pain (myalgia), joint pain (arthralgia), headache, and PEM.

### Data Visualization

Unless otherwise specified, all plots were generated in R with ggplot2. Volcano plots were made using the EnhancedVolcano package (Blighe et al., 2022).

## RESULTS

### Study Design

A cohort of 17 female sedentary control subjects and 18 females diagnosed with ME/CFS were subjected to a maximal cardiopulmonary exercise test (CPET) on a stationary bicycle. Blood was drawn from all subjects at baseline (0h), and at 15 minutes and 24 hours after the CPET. Extracellular vesicles (EVs) were isolated from the separated plasma and characterized using Nanoparticle Tracking Analysis (NTA) and Transmission Electron Microscopy (TEM). EV cargo was analyzed using nano liquid chromatography tandem mass spectrometry (nanoLC-MS/MS) untargeted proteomics (Figure 1).

**Figure 1:**
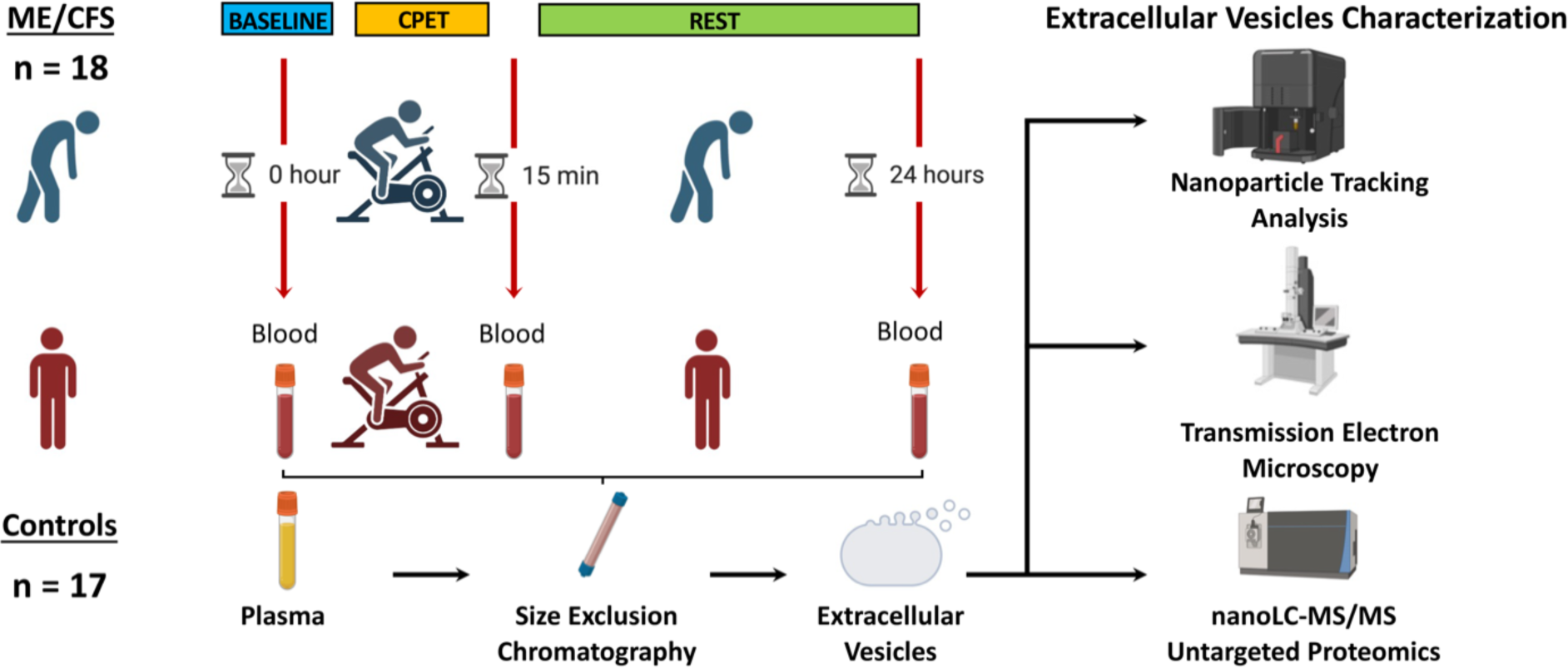
Study design including a Cardiopulmonary Exercise Test (CPET) and blood sample collection at baseline (0h), 15 minutes and 24 hours post-exercise, followed by extracellular vesicles (EVs) isolation by size exclusion chromatography and characterization of EVs by means of nanoparticle tracking analysis, transmission electron microscopy and nanoLC-MS/MS untargeted proteomics.

As shown in Table 1, subjects are age- and BMI-matched. Eighty-three percent of the ME/CFS patients report an acute, often flu-like illness that immediately preceded the onset of the disease, while 17% are unaware of an initiating event and consider their onset to be gradual (Table 1). The Bell Disability Scale scores clearly illustrates the condition of ME/CFS subjects versus controls, with a smaller score reflecting the lower functional level of patients compared to the higher score of functional controls. Both the Physical and Mental Component Summaries (PCS and MCS, respectively) derived from the SF-36 are, as expected, higher in the control group, indicating better health (Table 1).

### Characterization of EVs by Nanoparticle Tracking Analysis and Transmission Electron Microscopy

EVs were isolated from plasma by size exclusion chromatography at all three time points. EV size assessed by NTA did not significantly differ between healthy individuals and ME/CFS patients at any of the three time points (Supplementary Figure S2 and Supplementary Table S1). For EV concentrations, many differences were observed (Figure 2A and Supplementary Table S1). At baseline and 15 minutes post-exercise, the ME/CFS subjects had a higher concentration of circulating EVs compared to the control group (p = 0.0002 and p = 0.02, respectively). The mean EV concentrations in control subjects significantly increased by 1.4-fold at 15 minutes and 2.2-fold at 24 hours compared to baseline (p = 0.03 and p = 0.001 respectively, Figure 2A). The ME/CFS EV concentration was not significantly affected by exercise (Figure 2A).

**Figure 2:**
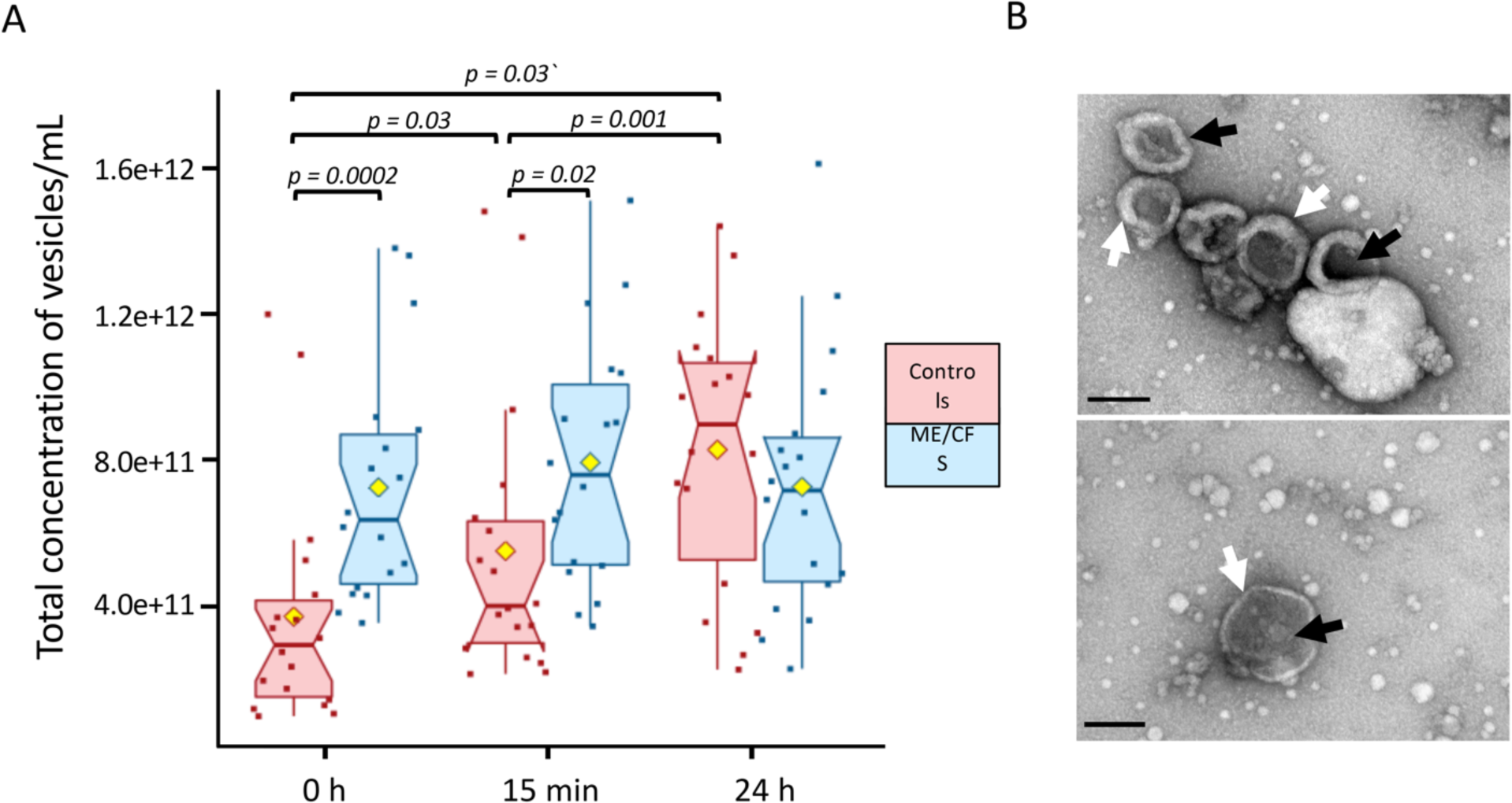
Nanoparticle Tracking Analysis and Transmission Electron Microscopy. (A) Total concentration of vesicles per mL of plasma in ME/CFS and healthy controls. The yellow dot represents the mean. (B) Representative TEM images obtained by negative staining, showing morphological structures of plasma-derived extracellular vesicles isolated by size exclusion chromatography. The black arrows highlight a few EV-like nanoparticles and the white arrows the characteristic lipid bilayer (scale bar: 200 nm). The non-parametric Wilcoxon signed-rank test was used to test the significance of differences (p < 0.05) between cases and controls.

Representative EVs were further analyzed by TEM confirming a population of vesicles with bilayered membranes and “saucer shape” features, which is a typical EV morphology (Figure 2B). The size distribution was consistent with the NTA analysis. However, smaller particles that possibly represent lipoprotein particles and a small number of large particles of 200–350 nm in diameter were also present in the samples analyzed.

### Characterization of EV cargo by quantitative proteomics

The EV protein content was further investigated by untargeted nano liquid chromatography tandem-mass spectrometry (nanoLC-MS/MS) TMT-based quantitative proteomics. A total of 862 proteins were detected (full list in Supplementary Table S2) and 301 were retained for further analysis after filtering out proteins with missing values for more than one third of subjects. 141 proteins had no missing data and the remaining missing values were imputed with random forest (RF, *missForest* R package, normalized root mean square error of 0.47). RF can handle non-linear data and outliers with no need for feature scaling and it was found to be the best method for imputing missing values in mass spectrometry data when the reason for the missingness is unknown (missing completely at random; MCAR) (Kokla et al., 2019; Wei et al., 2018). RF also has inherent feature selection, which makes it robust to noisy data (Breiman, 2001).

We cross-referenced our EV protein dataset with two manually curated EV proteome databases: Vesiclepedia (Kalra et al., 2012) and Exocarta (Keerthikumar et al., 2016). We observed over 94% overlap between our dataset and these databases (Figure 3), and we were able to detect a host of known EV markers such as tetraspanin CD9, vesicle trafficking-associated protein PDCD6IP (ALIX), heat shock proteins HSP90AA1 and HSPA8, and annexins ANXA3, ANXA5 and ANXA11.

**Figure 3:**
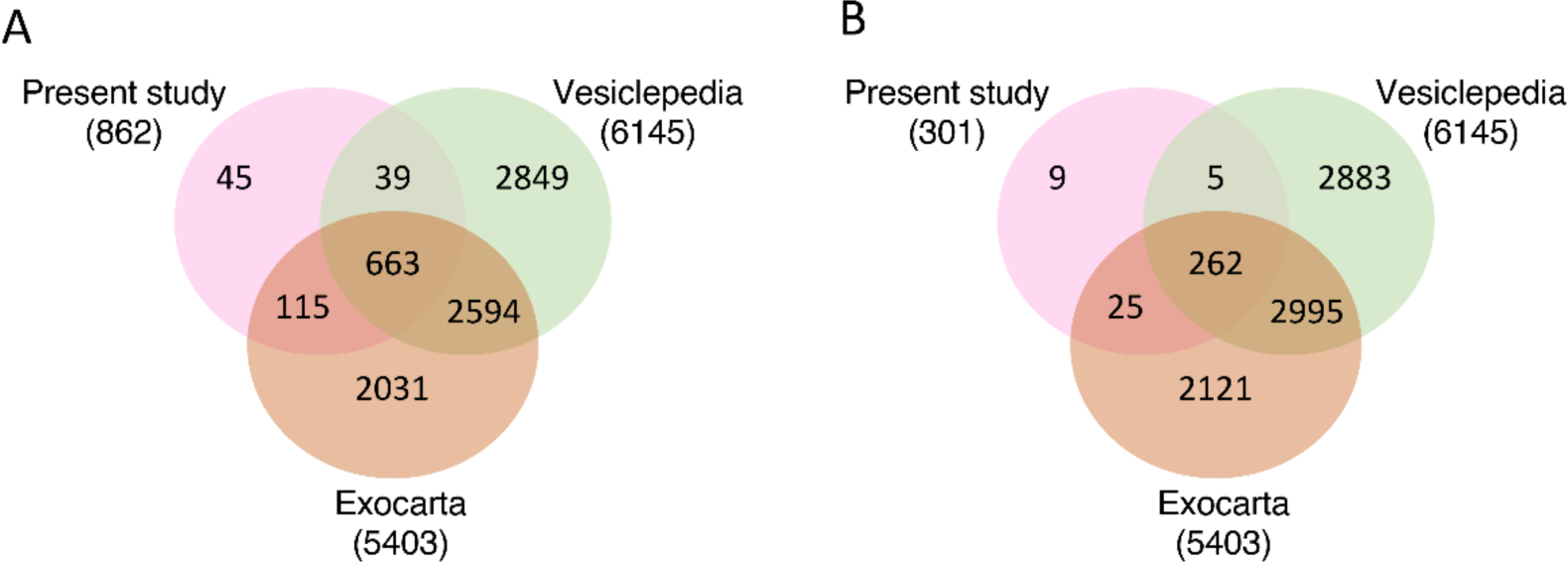
Deep proteomic coverage of the EV proteome. Comparison of proteomic coverage in our dataset versus manually curated vesicle databases Vesiclepedia and Exocarta of (A) all EV proteins identified and (B) after filtering out proteins with more than a third of missing values.

### Comparison of EV protein levels before and after exercise in ME/CFS patients vs. controls

Due to the experimental setup outlined in Supplementary Figure S1, samples could only be compared within each TMT 10-plex experiment. Differences in EV protein levels between ME/CFS and controls at each time point were analyzed using a bootstrapping strategy (details in Methods). Figure 4A shows the median Log2 fold change of ME/CFS subjects vs. controls and both the 95% and FDR-adjusted confidence intervals from 10,000 bootstrapped datasets. Protein levels are considered significantly different between groups if the adjusted confidence interval does not include 0. Those proteins are differentially abundant proteins (DAPs). Therefore, if the confidence interval is below 0, that DAP is significantly lower in ME/CFS compared to controls and if the confidence interval is above 0, then the DAP is significantly higher in the ME/CFS group.

**Figure 4:**
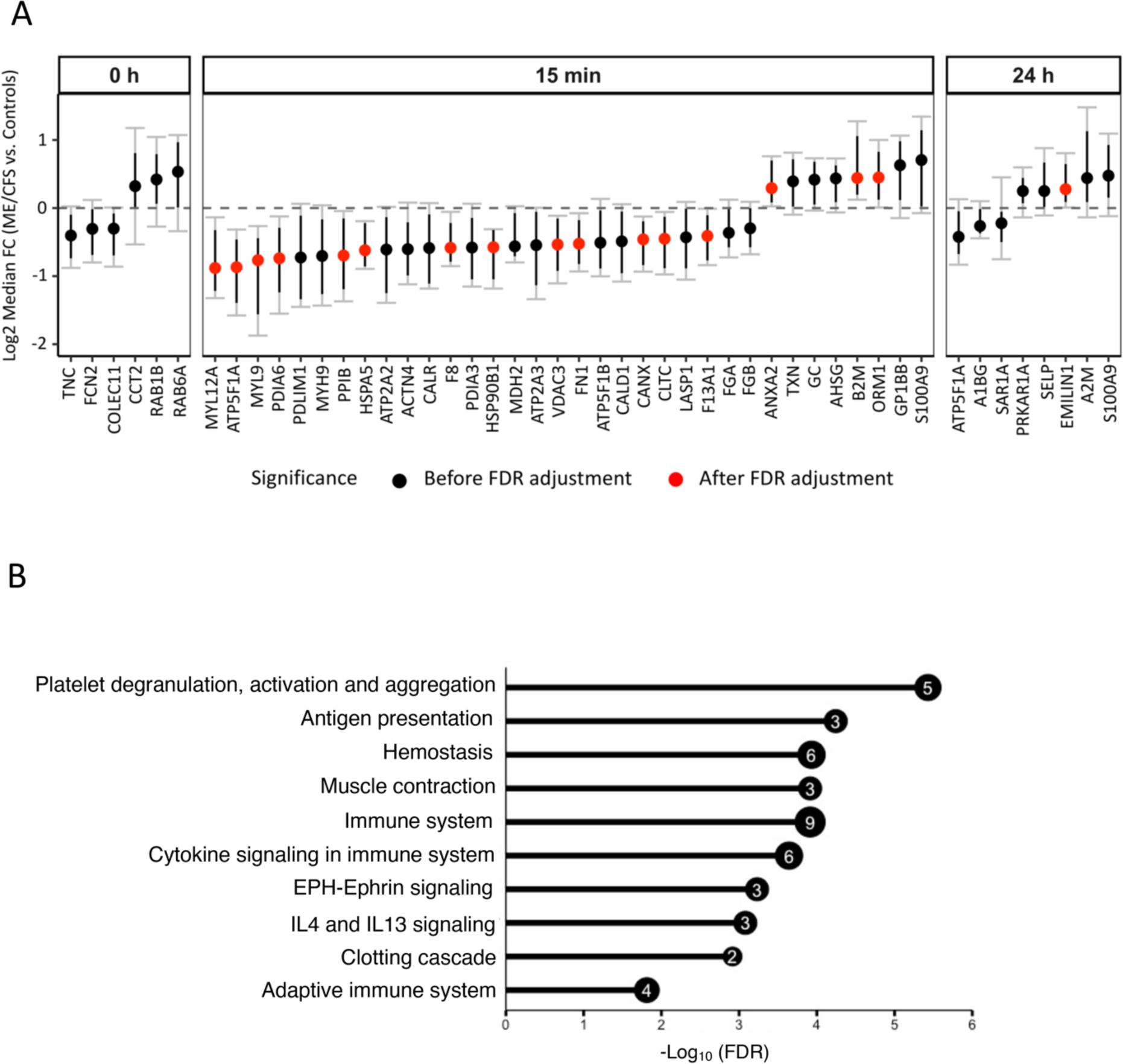
Differences in EV protein levels between ME/CFS and controls at each time point (A) and enrichment analysis (B). (A) The y-axis shows the Log2 fold change (ME/CFS vs. Controls) of 10,000 bootstrapped datasets at each time point. A median FC of 0 indicates no difference. Black dots show all proteins significant before FDR correction. Black bars show 95% confidence intervals (CI). Gray bars with caps show the false discovery rate adjusted CIs (with q < 0.1). A protein is significant after FDR correction (DAPs, red dots) if the adjusted CI does not include 0. (B) Bar plot showing - Log10(FDR) of the top 10 significant Reactome pathways (FDR <0.05) enriched for EV proteins that are significantly different (q < 0.1) between ME/CFS and controls 15 minutes post-exercise. The number inside the bubble shows the number of EV proteins in each pathway.

At baseline (0h), none of the six proteins significantly different before FDR correction remained significant after FDR-adjustment of the 95% confidence intervals (Figure 4A, black dots). 15 minutes post-CPET, 13 EV proteins had lower abundances and three had higher abundances in the ME/CFS group (Figure 4A, red dots, q < 0.1). Only one protein (EMILIN1) was significantly increased 24 hours post-exercise in ME/CFS patients (Figure 4A).

To gain functional insight into the proteomic cargo in purified EVs, we performed a Reactome pathway enrichment analysis on the DAPs 15 minutes post-exercise using *ShinyGO 0.76* (Ge et al., 2020) (Supplementary Table S3). The most significantly enriched Reactome pathways are illustrated in Figure 4B. Among them, “platelet degranulation, activation and aggregation” was the most enriched pathway including the coagulation factors F8 and F13A1, fibronectin 1 (FN1), and heat shock protein family A member 5 (HSPA5) which were decreased and orosomucoid 1 (ORM1) which was increased (Figure 4A). Other pathways related to blood physiology were enriched such as “hemostasis” and “clotting cascade”. The functional enrichment analysis also revealed that EVs at 15 minutes post-exercise contained many DAPs that were overrepresented in pathways related to the “immune system” (9 out of the 13 DAPs), “cytokine signaling”, “adaptive immune system” and “IL4-IL13 signaling”. Enriched pathways also included “muscle contraction” and “EPH-Ephrin signaling” which both include members of the myosin family, MYL12A and MYL9, that we found to be decreased in EVs from ME/CFS patients (Figure 4A).

### Ingenuity Pathway Analysis shows activated or inhibited canonical pathways in ME/CFS cases vs. controls

Functional analysis of the median Log2 fold change in ME/CFS patients vs. controls from the 10,000 bootstrapped datasets generated for each time point was performed using QIAGEN IPA. Enrichment analysis was performed using the list of 301 proteins analyzed in this study and the five most significantly enriched canonical pathways (marked with an asterisk in Figure 5) are “Acute phase response signaling”, “LXR/RXR activation”, “remodeling of epithelial adherens junctions”, “actin cytoskeleton signaling,” and “integrin signaling,”. The enrichment analysis was not time point dependent, as the datasets for all three time points include the same 301 proteins.

**Figure 5:**
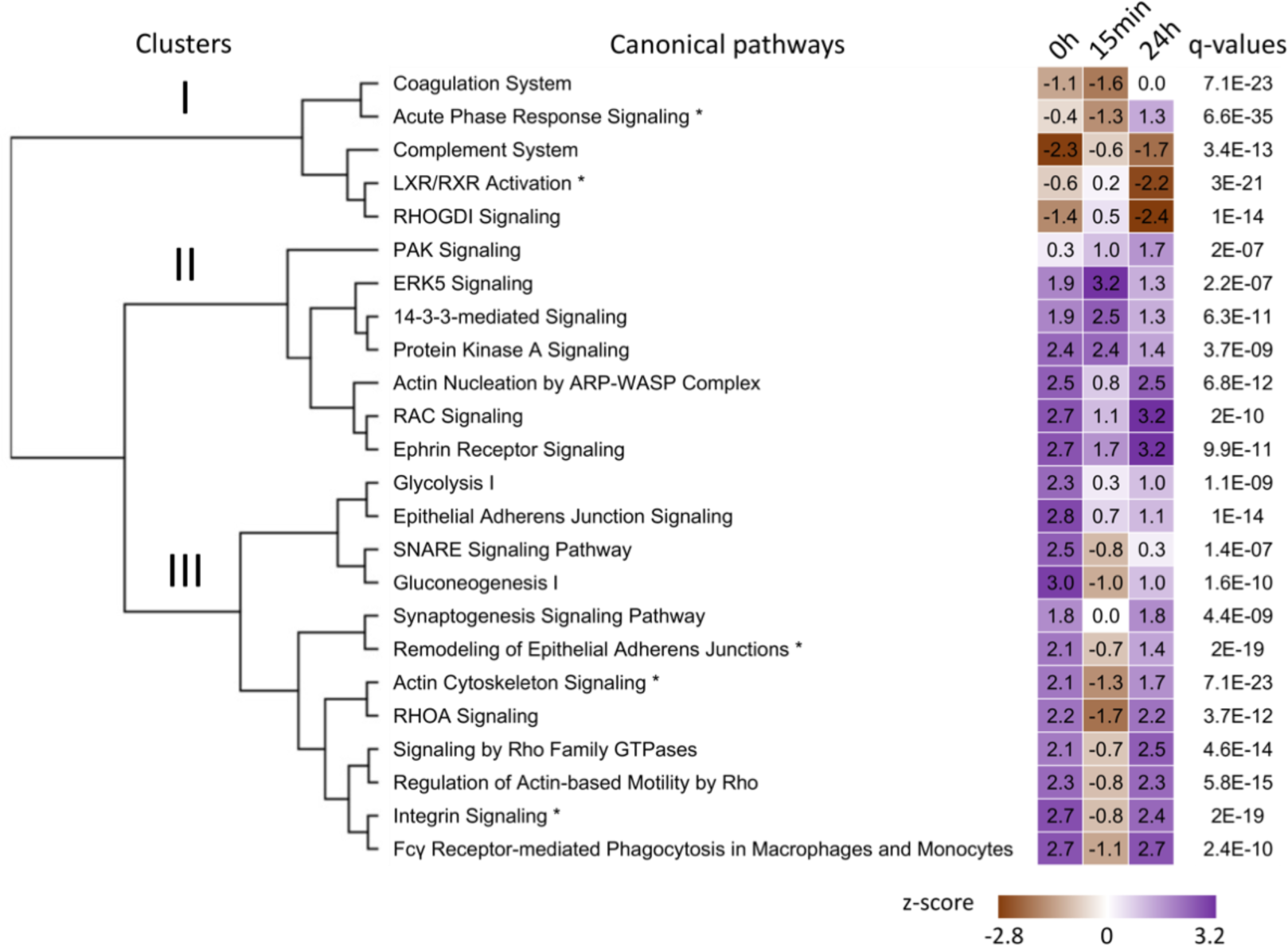
Output for the comparison analysis of the three timepoints using the QIAGEN IPA software. Top canonical pathways with z-scores were sorted by hierarchical clustering based on q-values (Fisher’s exact test followed by Benjamini-Hochberg correction). The top five canonical pathways are identified by an asterisk next to their name. The heatmap visualizes the z-scores of the canonical pathways at each time point. The sign of the calculated z-score reflects the overall predicted activation state of the biological function (<0: decreased, >0: increased). An absolute z-score of ≥ 2 is considered significant.

A comparison analysis between time points, using the median Log2 fold change in ME/CFS patients vs. controls for all 301 proteins, was also performed using QIAGEN IPA (Figure 5). In this analysis, a negative z-score reflects a predicted inhibition state of the canonical pathway in patients compared to controls, and a positive z-score indicates the pathway is activated in patients compared to controls. An absolute z-score of ≥ 2 indicates a significantly altered pathway in patients vs. controls at a specific time point.

Hierarchical clustering of the canonical pathways based on their z-scores at all three time points yielded three clusters (Figure 5). The five canonical pathways of cluster I were overall inhibited in patients compared to controls at all three time points (brown – negative z-scores). At baseline, only the “complement system” pathway was significantly inhibited in ME/CFS patients compared to controls (z-score = −2.3). While none were significantly altered at 15 minutes post-exercise, “LXR/RXR activation” and “RHOGDI signaling” were significantly inhibited 24 hours post-exercise (z-score = −2.2 and −2.4 respectively). The “complement system” and “LXR/RXR activation” are strongly involved in inflammation. The “coagulation system” is a complex process involving platelets and both the formation and degradation of clots requiring a finely balanced system for optimal function. “RHOGDI signaling” is involved in a multitude of biological processes including cell proliferation, apoptosis, cytoskeletal reorganization, and membrane trafficking.

Cluster II included seven canonical pathways that remained activated in patients compared to controls throughout the experimental protocol (purple – positive z-scores in Figure 5). Amongst them, the “14-3-3-mediated signaling” pathway was significantly activated in patients compared to controls at 15 minutes post-exercise (z-score = 2.5) but not 24 hours later.

Finally, cluster III included 12 canonical pathways that were initially activated in patients compared to controls and returned to that state 24 hours post-exercise, but were inhibited at 15 minutes post-exercise (Figure 5). All but “synaptogenesis signaling” were significantly activated at baseline in patients vs. controls (z-scores > 2). At 24 hours post-exercise, pathways related to the RHO GTPases family, “integrin signaling” and phagocytosis were activated in patients.

Apart from “glycolysis I” and “gluconeogenesis I” which are linked to energy metabolism and were significantly activated in ME/CFS patients vs. controls at baseline, most of the other canonical pathways are involved in actin cytoskeleton regulation for cell-cell communication including neural and immune cells.

### Changes in EV protein levels over time within groups

Changes in EV protein levels over time within the ME/CFS and control groups were assessed by calculating the within-subject fold changes for 15min vs. 0h, 24h vs. 0h, and 24h vs. 15min and the mean fold changes were compared to 1 using a student’s t-test. Amongst all comparisons performed we found significant results only at q < 0.15 for 15min vs. 0h and results are displayed as volcano plots in Figure 6A. A total of 63 and 187 DAPs were found in the ME/CFS and control groups, respectively, including 45 common to both (Venn diagram in Figure 6A). All 63 DAPs in the ME/CFS group had higher levels post-exercise (15min) compared to baseline (0h). In the control group, 178 DAPs had higher levels and nine had lower levels 15 minutes post-CPET (AGT, C4B, CLU, COLEC11, FCN2, MBL2, ORM2, PON1 and VSIG6). A Reactome pathway enrichment analysis revealed 276 and 298 pathways significantly enriched (q < 0.05) for the ME/CFS and control groups respectively (Supplementary Tables S4 and S5) with 189 pathways common to both groups.

**Figure 6:**
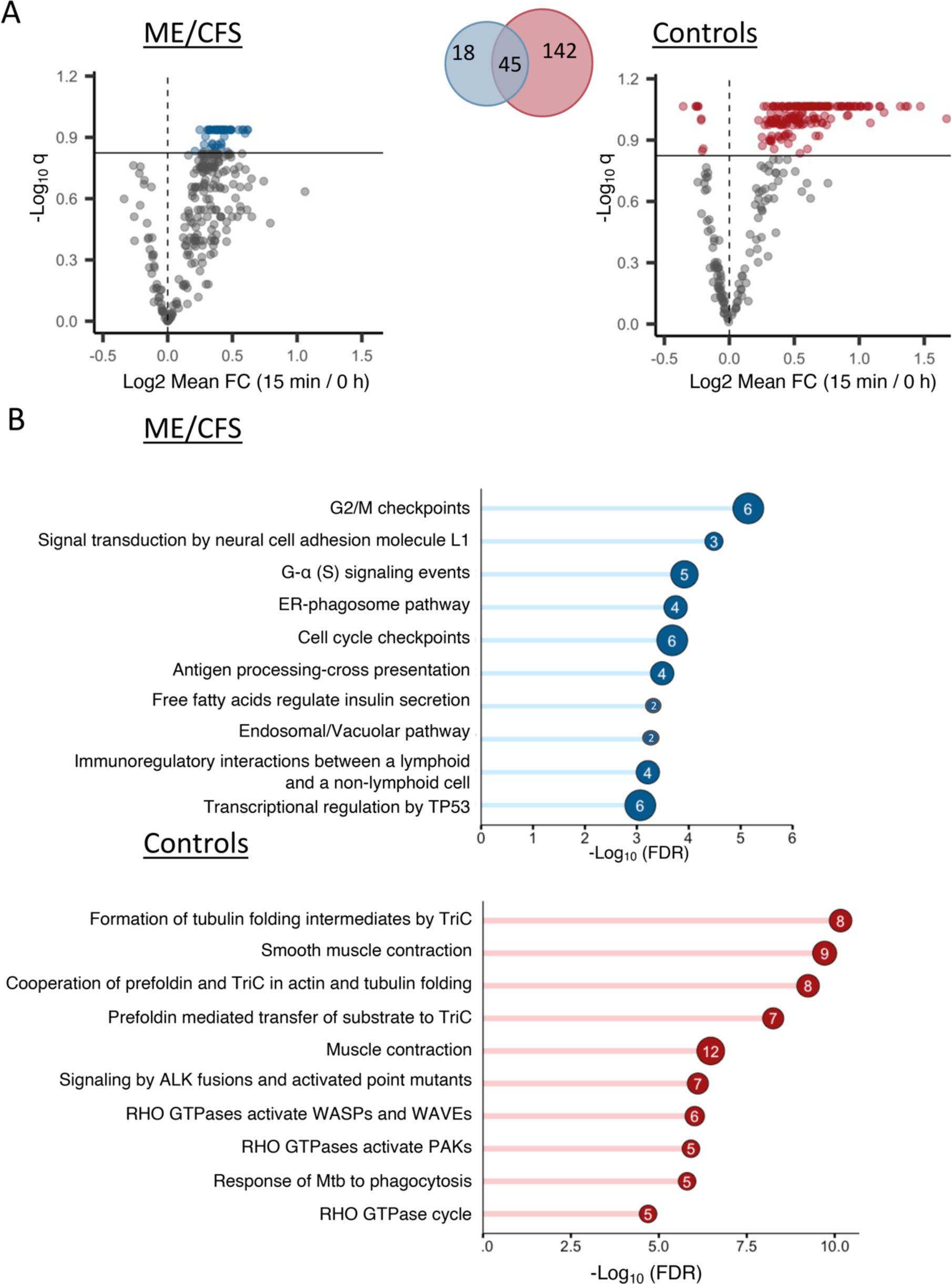
Changes in EV protein levels in ME/CFS and controls at 15 minutes post-exercise (A) and enrichment analyses (B). (A) Volcano plots showing significantly different EV protein levels in the ME/CFS and control groups at 15 minutes post-exercise compared to baseline. The x-axis shows the log2 mean fold change (15min vs 0h) and the y-axis shows the negative log of the q-value (a higher number represents increasing significance). The horizontal line shows the significance threshold q < 0.15 and colored dots are EV proteins that are significant at that threshold (gray dots are below the threshold). The Venn diagram is showing the number of significant EV proteins in both groups and their overlap. (B) Bar plots showing -Log10(FDR) of the top 10 significant Reactome pathways (FDR < 0.05) uniquely enriched in ME/CFS patients and controls. The number inside the bubble shows the number of EV proteins in each pathway.

We isolated pathways that were uniquely enriched in either the ME/CFS patients or the controls and found 85 enriched pathways unique to ME/CFS and 107 unique to controls (Supplementary Tables S6 and S7). Figure 6B shows the top ten unique Reactome pathways for each group. Within the ME/CFS group, the EV proteins with increased abundance include six members of the 14-3-3 protein family (YWHAE, YWHAQ, YWHAB, YWHAZ, YWHAG, YWHAH) that were significantly and uniquely associated with pathways related to “cell cycle checkpoints, or transcriptional regulation by TP53” and three EV proteins were integrins (ITGB1, ITGB3 and ITGA2B) associated with the “signal transduction by neural cell adhesion molecule L1” pathway. In the control group, EV proteins were overrepresented in pathways related to “actin and tubulin folding” (including members of the chaperonin-containing TCP1 ring complex TRiC (CCT2, CCT5, CCT6A, CCT8, TUBB, TUBB4B, TUBA4A, TUBA8)), “muscle contraction” (MYL6, MYL9, MYL12A, MYLK, TPM4, TMOD3), and “Rho GTPases”.

### Tissue and cell type enrichment analysis

We then performed tissue enrichment analysis using *Enrichr* (Chen et al., 2013) to explore which tissue and cell types typically express the proteins that have changing levels in EVs in ME/CFS patients or controls at 15 minutes post-exercise compared to baseline. The 187 DAPs in controls and the 63 DAPs in ME/CFS patients (q < 0.15) were compared to three databases containing reference gene and protein expression for different cell and tissue types (Human Gene Atlas, ARCHS4 Tissues, and HuBMAP ASCT+B augmented with RNAseq co-expression). Significant enrichment was determined using Fisher’s exact test and BH FDR correction (significance threshold q < 0.15). While we cannot determine the source of the EVs in the plasma of patients and controls, this analysis links the protein cargo of EVs to tissue and cell types in which they may have originated and reveals how the associated tissues and cell types are different in ME/CFS patients and controls after exercise.

We isolated cell and tissue terms that were uniquely enriched in either ME/CFS patients or controls and grouped them into broader physiological categories (see Supplemental Tables S8, S9 and S10 for the original terms and to which databases they belong). Sankey networks showing the links between proteins and enriched tissue types for controls and ME/CFS patients are shown in Figure 7A and B. Although there were 45 proteins significantly increased in abundance 15 minutes post-exercise in both ME/CFS patients and controls (Venn diagram, Figure 6A), only three of these proteins are contributing to the uniquely enriched tissue terms: TPI1, GNAI2, and PLEK. The broad categories that we found enriched in the controls but not in the patients include myoblasts, smooth muscle, kidney, and heart tissue. The only category that we identified to be exclusively enriched in the ME/CFS patients was lung tissue. Most of the 81 proteins in the control network are associated with either myoblasts, smooth muscle, or both. Muscle tissue terms were not significantly enriched in the ME/CFS patients, indicating that systemic signaling from muscle tissue through EVs post-exercise could be disrupted in ME/CFS patients and contributing to their pathophysiological response to exercise.

**Figure 7:**
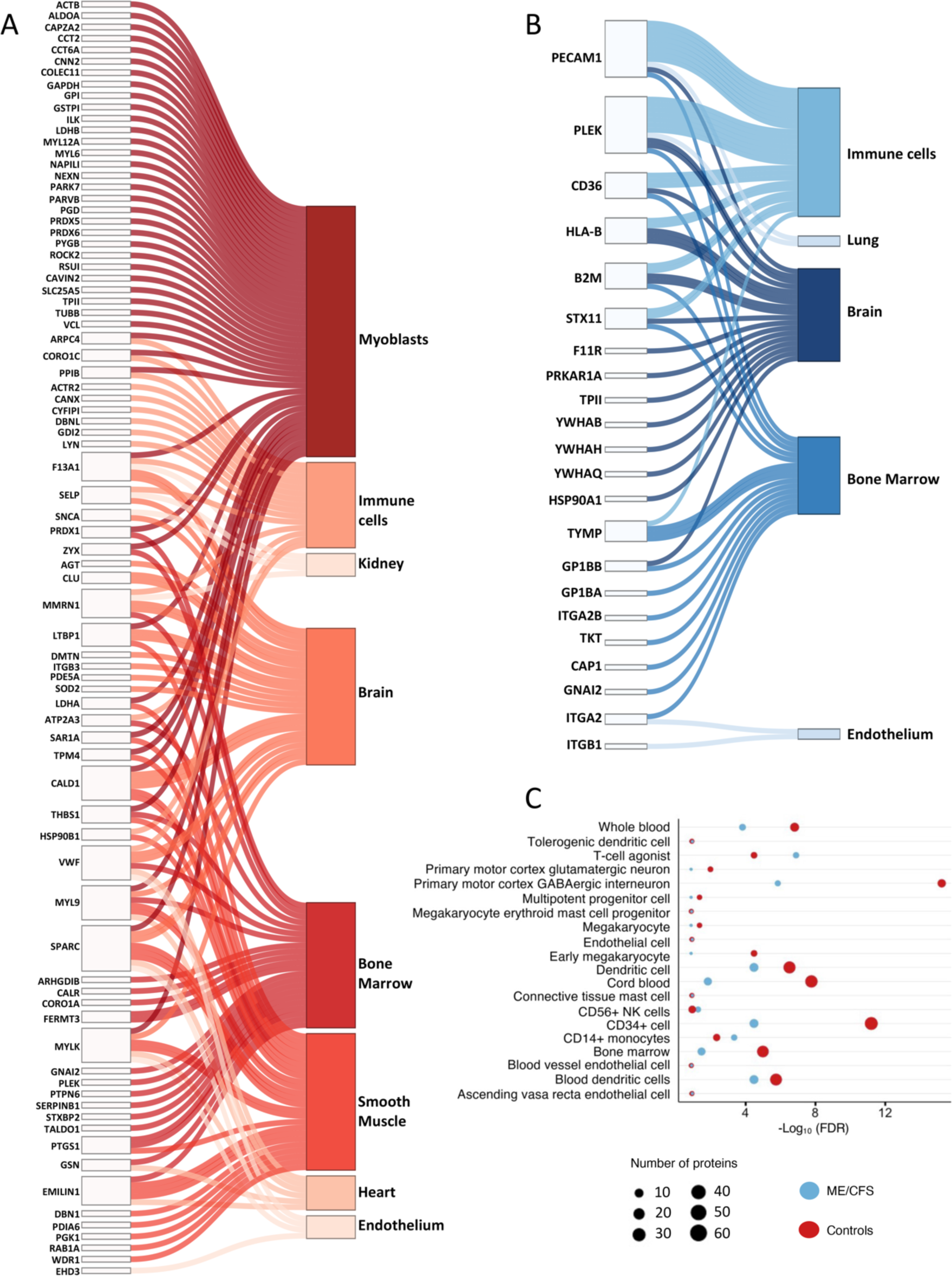
Tissue and cell type enrichment analysis. Sankey network diagrams showing 81 and 22 EV proteins contributing to uniquely enriched tissue type terms in controls (A) and in ME/CFS patients (B), respectively. Each line in the Sankey network shows one connection between a protein and an enriched tissue or cell type term. (C) Bubble plot with terms that are commonly enriched in both ME/CFS patients and controls, as well as their significance (-Log10 (FDR)) and the number of proteins overlapping with each reference gene/protein set.

While the categories immune cells, brain, bone marrow, and endothelium appear in both networks, the terms within those categories are unique to either patients or controls. Within immune cells, BDCA4+ Dendritic Cells and Tissue Resident Mucosal Type Mast Cells (id:605) were enriched only in controls. Despite a much smaller set of proteins changing significantly after exercise, the increased DAPs found in ME/CFS patients are significantly and uniquely associated with substantially more immune cell types, including Mast Cell (id:394), CD4 T Cell Memory (id:90), Non-classical Monocyte (id:475), Interstitial Macrophage (id: 348), Macrophage-Resident (id:388), Basophil (id:66), and multiple types of dendritic cells (Supplementary Table S8). Figure 7C shows the terms that are commonly enriched in both ME/CFS patients and controls, as well as their significance and the number of proteins overlapping with each reference gene/protein set. Within immune cells, the proteins changing in response to exercise in both patients and controls are associated with CD14+ Monocytes, CD56+ NK cells, Dendritic Cells, T-cell agonist (id:580), Connective Tissue Mast Cell (id: 133), and Tolerogenic Dendritic Cells (Tdc) (id:606). Many of the common terms are related to blood, bone marrow, and endothelium.

Within the brain category, only ME/CFS patients have changes in EV proteins that are expressed in the Subthalamic Nucleus and Amygdala which are both part of the basal ganglia of the brain (Chaudhuri & Behan, 2000). Since EVs can cross the blood-brain barrier, it is possible that EVs carrying these proteins originated in these parts of the brain. However, while these proteins are enriched in brain tissue, they are also expressed in other tissues (Thul & Lindskog, 2018), and we do not have information about which proteins are co-localized in specific EVs.

### EV proteins are changing differently over time in ME/CFS patients vs. controls in response to exercise

We used a similar bootstrapping approach to that described previously to compare the within-subject protein fold changes over time in ME/CFS patients vs. controls. Because the median fold change represents the difference between the change over time in ME/CFS vs. controls, but not the direction of that change within each group (red dots in Figure 8), we are providing the box plots at all three time point comparisons (15min/0h, 24h/0h, and 24h/15min, Supplementary Figure S3).

**Figure 8:**
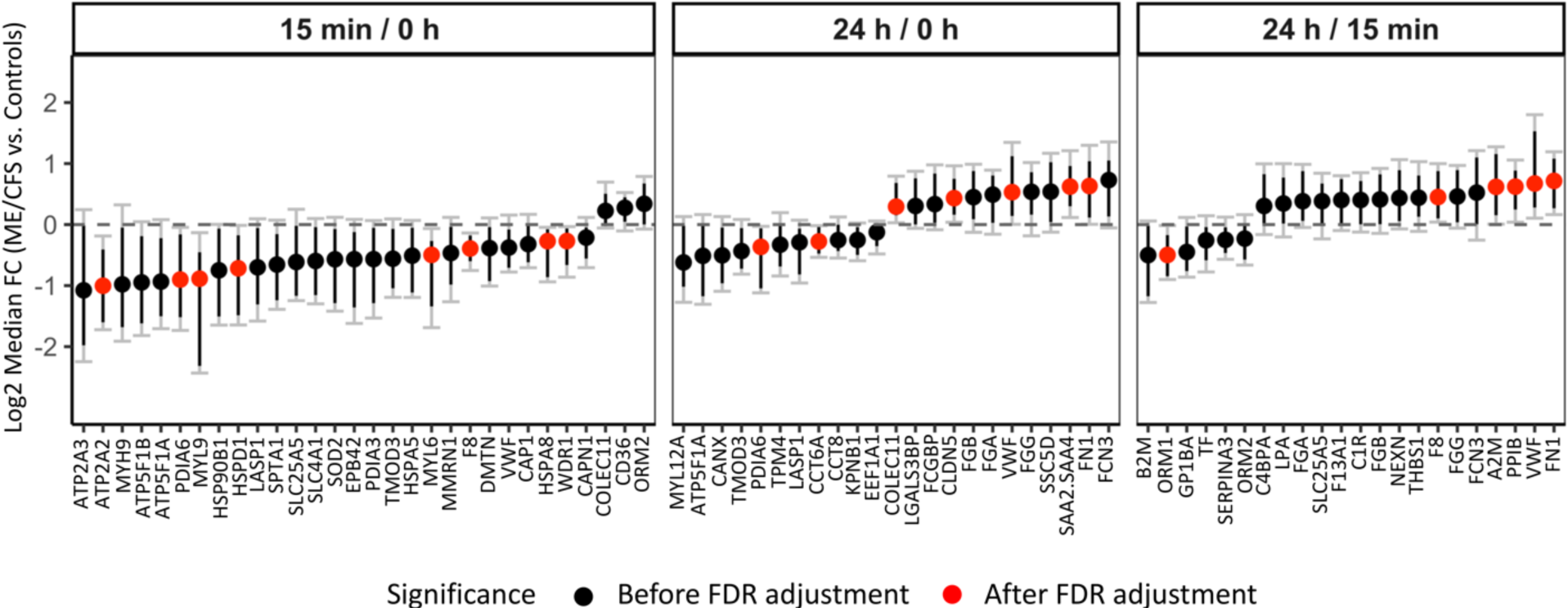
Changes in EV protein levels between ME/CFS and controls over time. The y-axis shows the median Log2 fold change (ME/CFS vs Controls) of 10,000 bootstrapped datasets for each time point comparison. A median FC of 0 indicates no difference. Black dots show all proteins significant before FDR correction. Black bars show 95% confidence intervals (CI). Gray bars with caps show the false discovery rate adjusted CIs (with q < 0.1). A protein ratio is significant after FDR correction (red dots) if the adjusted CI does not include 0.

After FDR correction several EV proteins were changing significantly differently in ME/CFS patients vs. controls in response to exercise (red dots, Figure 8). Eight EV proteins had lower 15min/0h ratios in ME/CFS patients vs. controls. Three of these proteins (ATP2A2, MYL6 and MYL9) are related to muscle physiology. ATP2A2 is an enzyme that catalyzes the hydrolysis of ATP and is involved in the regulation of the contraction/relaxation cycle of skeletal muscle. The myosin light-chain elements, MYL6 and MYL9, regulate the mechano-enzymatic function of myosin, i.e., the force produced during muscular cross-bridge cycles. PDIA6, HSPD1 and HSPA8 are chaperones implicated in a wide variety of functions including protection of the proteome from stress, proteolysis, folding, and transport of newly synthesized polypeptides, and play a pivotal role in the protein quality control system.

When comparing the change from baseline to 24 hours post-CPET (red dots, Figure 8), chaperone PDIA6 again had lower 24h/0h ratios in ME/CFS patients vs. controls. The 24h/0h ratio for CCT6A, a component of the chaperonin-containing TCP1 ring complex TRiC, was also lower while collectin COLEC11, claudin CLDN5, von Willebrand factor (VWF), the acute phase reactant serum amyloid read through (SAA2.SAA4), and fibronectin (FN1) ratios were higher in ME/CFS patients. Finally, when comparing 24 hours to 15 minutes post-exercise, only one protein had a lower 24h/15min ratio in ME/CFS patients (orosomucoid ORM1) and four were higher in patients including coagulation factor F8, alpha-macroglobulin A2M, PPIB, VWF and FN1.

### Changes in EV protein levels post-exercise strongly correlate with ME/CFS symptoms and disease severity metrics

For this analysis, Spearman’s correlation coefficients for the within-subject fold changes of the 301 EV proteins and selected clinical parameters were calculated. Bootstrapped 95% confidence intervals around the correlation coefficients from 2,000 bootstrapped datasets and associated p-values (null hypothesis that the correlation coefficient is 0) were calculated using the *bootcorci* R package, and q-values were obtained using the BH-FDR correction procedure. We define significant and strong correlations as those with |R| > 0.7 and q < 0.1. For the specific symptom severity (SSS) scores, each subject in both the ME/CFS and control groups was asked to rate their symptoms on a scale of 0-10 (0 = not present and 10 = very severe) at three different times: 1) on average over the past month (Initial), 2) on the morning of the CPET (0h) and 3) 24 hours after the CPET (24h).

The change in EV protein levels from 0h to 24h in ME/CFS patients had the greatest number of significant correlations at q < 0.1 (878 significant correlations; 648 R < 0 and 230 R > 0, Table 2, Supplementary Table S13). 69 of the negative correlations and 15 of the positive correlations also had |R| > 0.7. We found 72 significant correlations of clinical data with the proteins’ 15min/0h fold changes and 115 significant correlations for the 24h/15min fold changes (q < 0.1, Table 2).

**Table 2:** Number of significant correlations with clinical parameters (q < 0.1)

|  |  | R > 0 (> 0.7) | R < 0 (< -0.7) | Total |
| --- | --- | --- | --- | --- |
| Protein Fold Change | 15min/0h | 43 (8) | 29 (8) | 72 |
|  | 24h/0h | 230 (15) | 648 (69) | 878 |
|  | 24h/15min | 8 (2) | 107 (8) | 115 |

In parentheses are the number of correlations with $|R| > 0.7$ .

Two out of the eight SSS scores, the initial survey of muscle tenderness and pain (myalgia) and the initial survey of joint pain (arthralgia), had significant and strong correlations with the change in several EV protein levels from baseline to 24h. Figure 9A shows volcano plots of all correlation coefficients for those two symptoms. The proteins shown in dark blue on the volcano plots are those with Spearman |R| > 0.7 and q < 0.1 (32 and 34 proteins for myalgia and arthralgia, respectively, with eleven common to both symptoms, Figure 9A, Venn diagram). For both symptoms, only clusterin (CLU) had a strong positive correlation.

**Figure 9:**
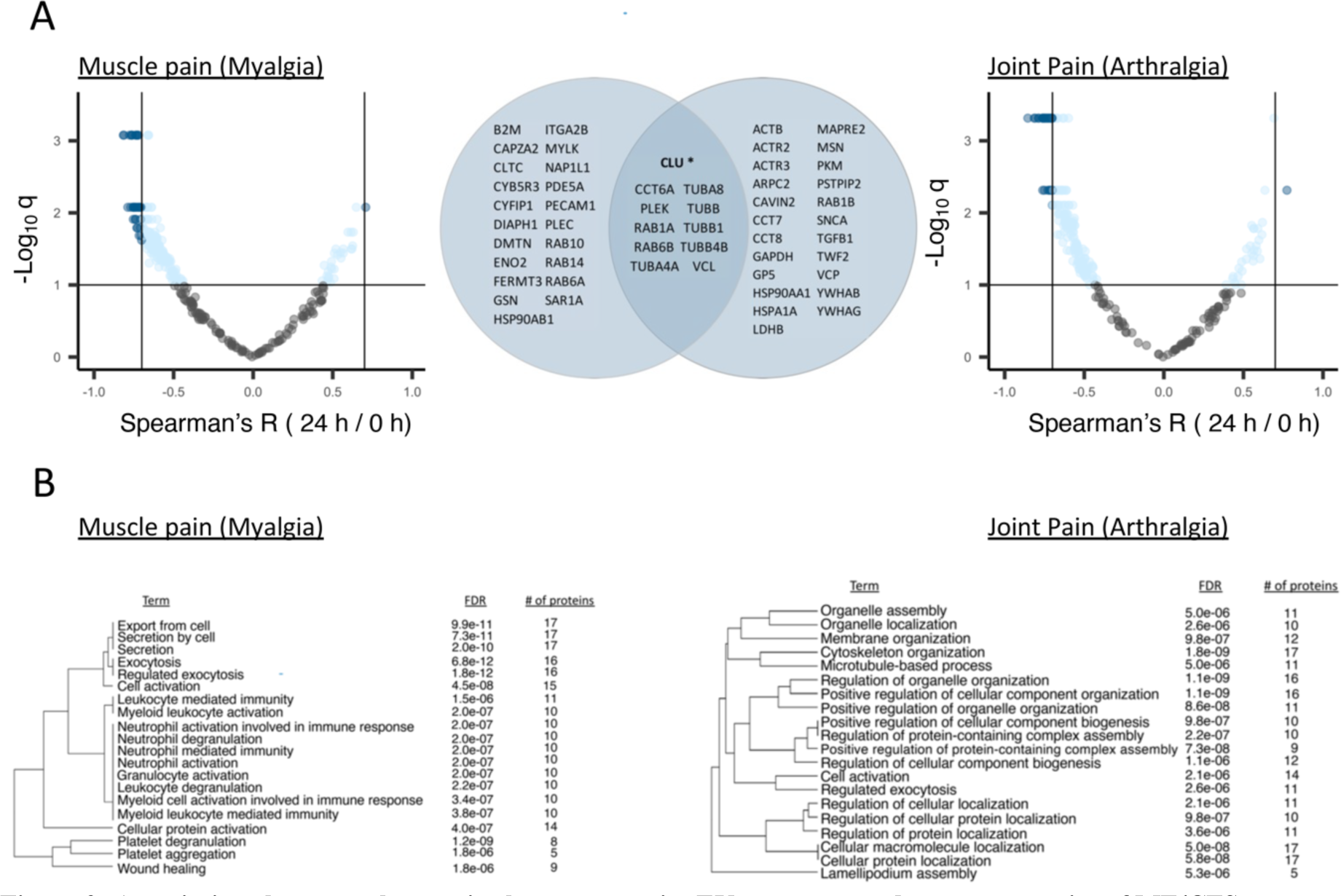
Associations between changes in the post-exercise EV proteome and average severity of ME/CFS symptoms myalgia and arthralgia. (A) Volcano plots showing correlations between the initial specific symptom severity (SSS) scores of muscle tenderness and pain (myalgia) and joint pain (arthralgia) and within-subject 24h/0h ratios for each protein. For initial SSS scores, subjects were asked to rate their average symptom severity over the last month. The x-axis shows the Spearman’s correlation coefficient R, with vertical solid lines indicating R = - 0.7 and R = 0.7. The y-axis shows the negative log of the q-value, with horizontal solid line indicating q < 0.1. The dark blue dots show proteins with |R| > 0.7 and q < 0.1, and light blue dots show the remaining proteins with q < 0.1. The Venn diagram shows the 32 and 34 proteins correlating at |R| > 0.7 and q < 0.1 with myalgia and arthralgia, respectively, with eleven being common to both. CLU* is the only protein with a positive correlation. (B) Pathway enrichment analysis of the 32 and 34 proteins displayed as hierarchical clustering dendrograms with the top 20 enriched pathways. Pathways with many shared proteins are clustered together. FDR and number of proteins belonging to each term are shown.

Pathway enrichment analysis was performed on these 32 and 34 strongly correlated proteins (|R| > 0.7 and q < 0.1) and displayed as hierarchical clustering dendrograms in Figure 9B. For the 32 proteins correlating with myalgia, there was an enrichment of pathways related to cell export, “leukocyte mediated immunity”, neutrophil activities, and platelet degranulation and aggregation (Figure 9B). For the 34 proteins correlating with arthralgia, pathways related to “cytoskeleton organization”, cellular component assembly and biogenesis, and “regulation of cellular protein localization” were the most significantly enriched (Figure 9B). Several of these proteins also contributed to tissue enrichment terms that were unique in ME/CFS patients (Figure 7B), including B2M, ITGA2B, PECAM1, and PLEK for myalgia and PLEK and YWHAB for arthralgia.

For all SSS scores on the days that blood was collected, we calculated the change in symptom severity post-exercise as the difference in SSS from 0h to 24h post-exercise (ΔSSS). Thus, a positive ΔSSS indicates a worsening of that symptom 24 hours post-exercise and a negative ΔSSS indicates that symptom improved 24 hours post-exercise. The 24h/0h ratios for seven proteins (CALM2, DBN1, GSTP1, HSPB1, THBS1, TMOD3 and TPM4) had strong and significant positive correlations to the change in myalgia after exercise in ME/CFS patients (Figure 10A), such that when the protein level increased from 0 to 24 hours (Log2FC > 0), myalgia also increased (Δ > 0), and when the protein level decreased post-exercise, myalgia was also decreased. Additionally, the change in CAPZB levels from 0h to 15min post-exercise also positively correlated with the change in myalgia (Figure 10B). Tropomodulin (TMOD3), tropomyosin (TPM4), and capping actin protein of muscle z-line subunit beta (CAPZB) are involved in muscle contraction. All of these correlations were absent in controls.

**Figure 10:**
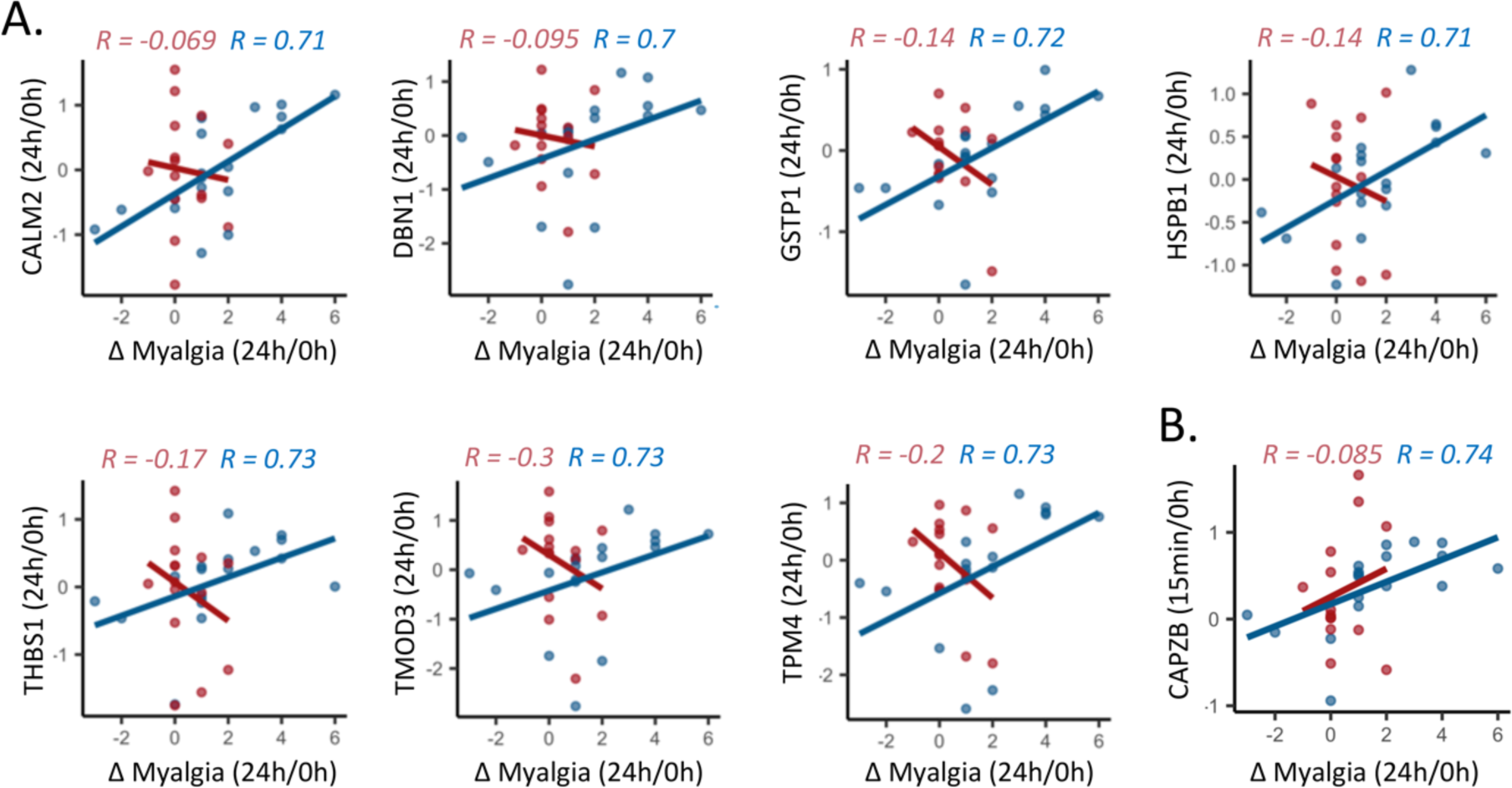
Correlations between the change in myalgia from baseline to 24 hours post-exercise and the within-subject 24h/0h ratios (A) or within-subject 15min/0h ratios (B). Each dot is one subject. The lines are linear regression lines for each group, ME/CFS or control. Spearman’s R is shown on each plot for controls (red) and ME/CFS patients (blue). All correlations presented are significant in ME/CFS patients (q < 0.1).

Two core symptoms of ME/CFS are post-exertional malaise (PEM) and fatigue. The SSS scores for PEM correlated with the change in several proteins post-exercise. In particular, the 15min/0h ratio of the two fibrinogen chain proteins FGA and FGB positively correlated with PEM that subjects were experiencing 24 hours post-exercise (Figure 11A). The 24h/0h ratio of nexilin f actin binding protein (NEXN) levels in ME/CFS patients correlated with ΔPEM such that an increase in NEXN correlated with a larger increase in PEM after exercise (Figure 11A). We found the 15min/0h ratios of two proteins, fibrinogen gamma chain (FGG) and clusterin (CLU), positively correlated with worse fatigue on the morning of the CPET (Figure 11B).

**Figure 11:**
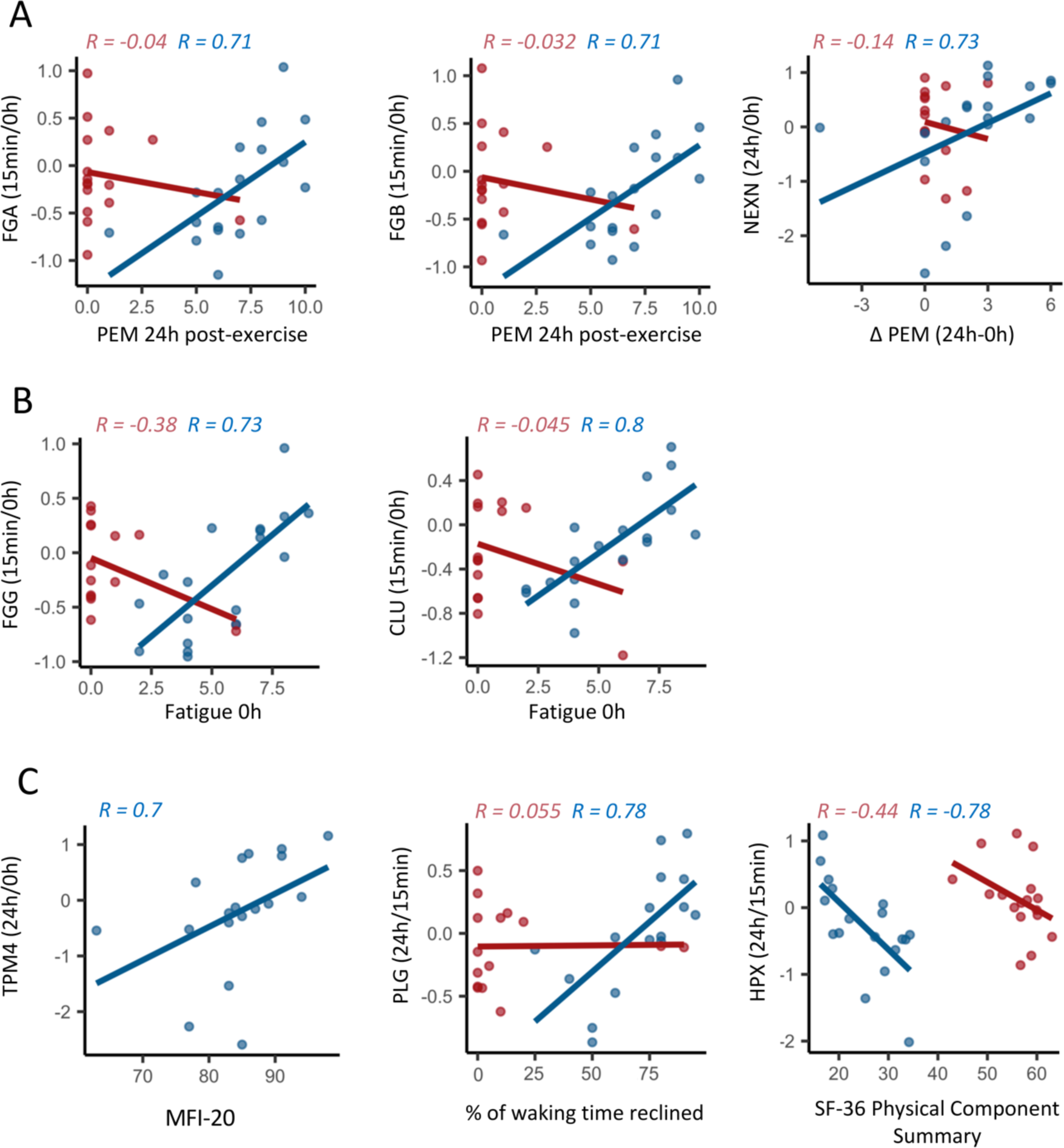
Associations between changes in the post-exercise EV proteome and (A) post-exertional malaise severity, (B) fatigue severity the day of exercise and (C) metrics of overall disease severity, including the MFI-20, the percentage of waking time spent reclined and the SF-36 Physical Component Summary. Each dot is one subject. The lines are linear regression lines and Spearman’s R are shown on each plot for controls (red) and ME/CFS patients (blue). All correlations presented are significant in ME/CFS patients (q < 0.1).

In addition to the SSS survey data, we also looked at correlations to survey data that are indicators of overall disease severity (Figure 11C). We found that the 24h/0h ratio of TPM4 positively correlated with the multidimensional fatigue inventory 20 (MFI-20) total score. For the MFI-20, a higher score corresponds to worse fatigue. The 24h/15min ratio of plasminogen (PLG) positively correlated with the percentage of waking time spent reclined or lying down (Figure 11C). A higher percentage of waking time spent reclined indicates worse overall ME/CFS severity. The 24h/15min ratio of hemopexin (HPX) negatively correlated with the SF-36 physical component summary (PCS) (Figure 11C), where higher scores indicate better health.

## DISCUSSION

Overall, this study characterized changes in the EV proteome of healthy sedentary control females in response to a single bout of exercise, while also demonstrating highly disrupted EV signaling post-exercise in ME/CFS patients. This is evidenced by: 1) fewer differentially abundant proteins (DAPs) in EVs 15 minutes post-exercise in patients compared to controls (Figure 4A); 2) 63 EV proteins increasing due to exercise in ME/CFS patients whereas 178 EV proteins increased in controls (only 45 overlapping) (Figure 6A); 3) a delayed increase in abundance of several EV proteins in response to exercise in ME/CFS patients compared to controls (Figure 8, Supplementary Figure S3). Thus, ME/CFS patients exhibit altered EV signaling dynamics and failure to mount an adequate response to exercise at the molecular level.

Multiple enrichment analyses were performed to enhance our understanding of the impact of this dysregulation of the EV proteome post-exercise on protein pathways and what tissues and cell types express these proteins. Taken together, these results show that the DAPs are involved in many pathways and systems, including platelet functions, muscle contraction (both smooth and skeletal muscle), cytoskeletal protein functions, the immune system, and brain signaling.

Several phenomena may contribute to this dysregulated signaling: 1) the tissue and cellular sources of the EVs may differ; 2) the EV cargo may differ between cases and controls, even when released from the same source; 3) cases and controls may differ in EV targets and/or uptake by destination cells and tissues.

Finally, we found strong and significant correlations between the change in EV protein levels post-exercise in ME/CFS patients and clinical metrics (myalgia, arthralgia, post-exertional malaise (PEM), fatigue, % of waking time spent reclined, and SF-36 physical component summary) (Figures 9-11). This suggests that the profound disruption of EV signaling post-exercise may contribute to the inability of ME/CFS patients to recover from exertion.

### EV concentration is differently affected by exercise in ME/CFS patients and controls

Plasma EVs were isolated from ME/CFS subjects and healthy controls and quantified by NTA. Our study revealed a significantly higher concentration of circulating EVs in individuals with ME/CFS compared to healthy controls at baseline and 15 minutes post-CPET (Figure 2), which supports recent findings previously reported in ME/CFS (Almenar-Pérez et al., 2020; Castro-Marrero et al., 2018; Eguchi et al., 2020; Giloteaux et al., 2023; Giloteaux et al., 2020). Similar observations have also been documented in other conditions, including Alzheimer’s disease (Rajendran et al., 2006) and cerebrovascular diseases (Jung et al., 2009).

In the control cohort, there was a significant increase in the total concentration of EVs 15 minutes and 24 hours post-exercise as compared to baseline, while no changes were detected in the ME/CFS group. Multiple studies conducted in both humans and rodents have documented a rapid rise in EVs following a single bout of physical exercise, leading to changes in EV plasma concentrations, markers, and contents (Bei et al., 2017; Brahmer et al., 2019; Oliveira Jr et al., 2018; Whitham et al., 2018). In a study with 17 healthy and 15 prediabetic men performing an acute cycling bout, a significant increase of the total concentration of vesicles in plasma was found exclusively in the healthy group during exercise from baseline to 30 minutes; however, in the prediabetic group, the total concentration of vesicles in the plasma was not affected by exercise (Warnier et al., 2022). Thus, elevated EV concentration relative to healthy controls at baseline and post-exercise may be explained by the inflammatory state characteristic of the disease.

### Large-scale increases in EV protein levels in healthy sedentary females post-exercise

We found 62% of proteins analyzed to be differentially abundant at 15 minutes post-exercise in healthy control females, with the vast majority increased, while none were significantly altered 24 hours post-exercise. This is consistent with a study characterizing the EV proteome of healthy males in response to a single bout of cycling exercise where differences were observed immediately post-exercise in 322 out of 1199 proteins analyzed, with most of them increased and only 3 DAPs four hours post-exercise (Whitham et al., 2018). It seems that in both females and males, the increase in EV protein abundance post-exercise is a transient phenomenon. The DAPs in EVs post-exercise in healthy sedentary females are expressed by many cell and tissue types, indicating a systemic, coordinated response to exertion by smooth and skeletal muscle tissue, the immune system, the brain, the kidney, as well as circulating cells and blood vessel endothelial cells (Figure 7A, C).

### Dysregulated coagulation processes in ME/CFS post-exercise

Throughout our analyses, we found many DAPs between ME/CFS patients and controls involved in coagulation processes. The coagulation cascade proteins factor VIII (F8), factor XIII A1 (F13A1) and fibronectin (FN1), were significantly decreased 15 minutes post-exercise in ME/CFS vs. controls (Figure 4A, B). F8 and FN1 also changed significantly over time in ME/CFS patients vs. controls (F8 at 15min/0h and 24h/15min; FN1 at 24h/0h and 24h/15min; Figure 8). Von Willebrand factor (VWF) had a similar expression pattern as FN1 (Figure 8, Supplementary Figure S3).

Changes from baseline to 15 minutes post-exercise in the fibrinogen chain proteins, FGA, FGB, and FGG, which are also part of the coagulation cascade, positively correlate with the ME/CFS symptoms fatigue and PEM post-exercise (Figure 11A, B). Additionally, the pathway analysis of the 32 proteins strongly correlating with general myalgia severity included “platelet degranulation” (8 proteins), “platelet aggregation” (5 proteins), and “wound healing” (9 proteins) (Figure 9B). Finally, the change in plasminogen (PLG) from 15 minutes post-exercise to 24h post-exercise (recovery) strongly correlated with the percentage of waking time spent reclining or lying down (a measure of overall disease severity), where a higher percentage of waking time spent reclining corresponds to more severe ME/CFS (Figure 11C).

Blood coagulation is enhanced in response to acute exercise in healthy individuals (Menzel & Hilberg, 2011) and in those with cardiovascular diseases (Mustonen & Lassila, 1997). This elevated coagulation potential is usually accompanied with increased fibrinolytic activity (Womack et al., 2003) to maintain hemostatic balance. Exercise-induced increases in F8 activity have been reported (Arai et al., 1990; Davis et al., 1976), and align with our results of increased F8 levels in control EVs 15 minutes post-exercise. Furthermore, muscle FN1 abundance was found to increase after moderate exercise (Mavropalias et al., 2023). Specific to EVs, Kobayashi and colleagues examined the EV proteome before and 30 minutes after high-intensity interval cycling in three healthy males and found significant changes in 20/558 proteins analyzed including increased abundance of FN1, VWF, FGA, FGB, and FGG (Kobayashi et al., 2021).

Post-exercise increases in F8, FGA, FGB, and FGG were also found in eleven healthy males (Whitham et al., 2018). Here, we found significant increases 15 minutes post-exercise for VWF and F8 in healthy females, but no significant changes in FN1, FGA, FGG, or FGB (Figure 6A). Also, increases in FGA and FGB in patients 15 minutes post-exercise correlated with worse PEM 24 hours post-exercise, and increases in FGG post-exercise correlated with worse fatigue the morning of the exercise test (Figure 11A, B).

EVs play a dual role in blood coagulation by promoting clot formation and initiating the conversion of plasminogen to plasmin, leading to fibrinolysis. We found that a larger increase in plasminogen abundance in EVs from 15 minutes post-exercise to 24 hours post-exercise was associated with worse overall ME/CFS severity (Figure 11C). This indicates a balance shift toward fibrinolysis during recovery which could be detrimental. EVs also have a transient hypercoagulable function and may play a role in the early phase of hemostasis after injury in trauma patients, with an increased concentration in the first 3 hours after injury (Rognes et al., 2021). The altered temporal profiles of clotting cascade factors in ME/CFS EVs post-exercise reveals a disruption in the hemostatic balance between clot formation and fibrinolysis.

Levels of coagulation-related EV proteins are also altered in other disease states. For instance, FGA, FGB, FGG, VWF, and F13 all had increased abundance in EVs of patients with idiopathic inflammatory myopathies compared to healthy controls (Meng et al., 2022). A recent proteomic analysis revealed F13A1, FGA, and FGB were all enriched in EVs from COVID-19 patients (Setua et al., 2022). FGB was also highly abundant in EVs of non-critical COVID-19 patients (Sur et al., 2021), while another study conducted on critical COVID-19 patients requiring mechanical ventilation found decreased EV fibrinogen components (Barberis et al., 2021).

Taken together, our results show that an altered hemostatic balance between coagulation and fibrinolysis may be contributing to PEM and exercise intolerance in ME/CFS patients. Other recent reports corroborate the importance of coagulation processes in ME/CFS. Dysregulation in platelet gene expression was found at 24 hours following maximal exercise (Ahmed et al., 2022). Likewise, an approximately 10-fold greater occurrence of fibrinaloid microclots was discovered in people with ME/CFS compared to healthy controls, which is clear evidence of hypercoagulability in ME/CFS even in the absence of an exercise challenge (Nunes et al., 2022). Considering the importance of EVs in maintaining hemostatic balance, future studies exploring the complex interplay between EVs, platelets, and the endothelium in ME/CFS patients are warranted.

### EV signaling relating to muscle contraction is impaired in ME/CFS

Our analyses also reveal impaired EV signaling in proteins involved in muscle contraction in ME/CFS patients, and some of these proteins were significantly associated with the symptoms myalgia (Figure 10) and fatigue (Figure 11C). Muscle contraction was in the top ten altered pathways at 15 minutes post-exercise in ME/CFS patients vs. controls (Figure 4B). The DAPs in controls 15 minutes post-exercise were associated with muscle and smooth muscle contraction pathways (Figure 6B), as well as myoblast and smooth muscle cell and tissue types (Figure 7A), while no muscle related tissue or protein pathway enrichment was found in patients. Myosin light chains MYL9 and MYL12A had decreased abundance in ME/CFS patients vs. controls 15 minutes post-exercise (Figure 4A). MYL9 and myosin light chain MYL6 were significantly increased post-exercise in EVs in controls but not in patients (Figure 6A) and this change over time was significantly different between groups (Figure 8). Indeed, in healthy controls, the protein with the largest mean increase 15 minutes post-exercise was MYL9 (3.2-fold) and MYL12A and MYL6 both showed greater than 2-fold increases (Supplementary Table S11). MYL12A, MYL6, and TPM4 were also found to be increased in EVs post-exercise in healthy males (Whitham et al., 2018).

Increases in EV levels of tropomyosin TPM4, tropomodulin TMOD3, and calmodulin CALM2 from baseline to 24 hours post-exercise were strongly correlated with higher levels of myalgia after exercise in the ME/CFS subjects (Figure 10). The 24h/0h ratio of TPM4 was also positively correlated with the MFI-20, which measures fatigue (Figure 11). Finally, the 24h/0h ratio of myosin light chain kinase (MYLK) was one of the many strong negative correlations with the average muscle pain rated by the patients over the last month (Figure 9). During exercise, myosin, tropomodulin, and tropomyosin all play crucial roles in muscle contraction and the regulation of muscle function in both smooth and skeletal muscle (Kishi et al., 1998).

The contribution of skeletal-muscle derived EVs to the post-exercise EV circulatory pool is not completely understood. Whitham et al. identified candidate myokines coming from skeletal muscle EVs by isolating EVs from both the femoral artery and femoral vein (Whitham et al., 2018). There is also strong *in vitro* evidence that myoblasts, myotubes, and skeletal muscle tissue explants release EVs (Estrada et al., 2022; Forterre et al., 2014). However, another study examining the surface markers of exercise-induced EVs in male athletes found no change in skeletal muscle markers and instead found increases in markers associated with lymphocytes, monocytes, platelets, endothelial cells and antigen presenting cells (Brahmer et al., 2019). More recently, Estrada and colleagues, using immunocapture of plasma EVs from transgenic mice expressing eGFP only in skeletal muscle, showed that ∼ 5% of CD9+ and CD81+ EVs were derived from skeletal muscle (Estrada et al., 2022).

The smooth muscle protein elastin microfibrillar interface protein 1 (EMILIN1) had increased abundance in ME/CFS patients vs. controls 24 hours post-CPET (Figure 4A). EMILIN1 is an ECM component that is abundantly expressed in elastin-rich tissues such as the blood vessels, skin, heart, and lung (Colombatti et al., 2000). EMILIN1 produced by vascular smooth muscle cells is a major regulator of resting blood pressure levels (Litteri et al., 2012). Interestingly, EMILIN1 is proteolyzed and secreted in small EVs as a mechanism to reduce its intracellular level (Amor López et al., 2021). Smooth muscle cells in the vasculature undergo a number of changes in response to acute exercise (Newcomer et al., 2011), secrete EVs *in vitro* and *in vivo* (Kapustin & Shanahan, 2016), and are responsive to EV signaling from vascular endothelial cells *in vitro* (Boyer et al., 2020). However, exercise-induced changes in EVs derived from smooth muscle cells have not been studied.

As of now, we are unable to determine the origin of the measured circulating EVs in this study. However, our analysis shows that a transient post-exercise increase in EV muscle-related proteins is part of the healthy response to exercise which is reduced in ME/CFS patients. The delayed increase in some of these EV proteins at 24 hours post-exercise in ME/CFS patients is associated with worse skeletal muscle tenderness and pain at the same time point (Figure 10).

### Changes in cytoskeletal proteins correlate with ME/CFS symptoms

We observed many correlations between changes in EV cytoskeletal proteins and ME/CFS symptom severity. Pathway analysis of the proteins correlating with arthralgia shows that these proteins are involved in cytoskeleton organization (17 proteins) and microtubule-based processes (11 proteins) (Figure 9). The 24h/0h ratios of tubulin proteins (TUBB, TUBA4A, TUBA8, TUBB1, TUBB4B) are negatively correlating with the general severity of both myalgia and arthralgia, so higher levels of the proteins 24 hours post-exercise compared to baseline correspond to lower levels of pain (Figure 9A). TUBA1B/A/C measured in plasma was recently included in a set of 20 proteins (15 measured in plasma and 5 measured in EVs) used to classify ME/CFS patients vs. controls with 86% accuracy (Giloteaux et al., 2023).

We observed an upregulation of tubulins (TUBB, TUBB4B, TUBA4A, TUBA8) and members of the TRiC complex (CCT2/5/6A/8) in EVs from controls 15 minutes post-exercise (Figure 6B). TUBB and CCT2 are both contributing to the enrichment of the myoblast cell type in controls (Figure 7). The 24h/0h ratio of CCT6A was significantly different between patients and controls (Figure 8, Supplementary Figure S3) and is negatively correlated with both myalgia and arthralgia in ME/CFS patients (Figure 9A). Microtubules are one of the principal components of the cytoskeletal system and are involved in muscle development and contraction, but they are also involved in maintenance of neuronal morphology and the formation of axonal and dendritic processes (Baas et al., 2016; Becker et al., 2020).

Changes 24 hours post-exercise in proteins related to actin polymerization, including actin beta (ACTB), actin-related protein 2 (ACTR2), and actin-related protein 3 (ACTR3) are also negatively correlating with arthralgia in ME/CFS patients (Figure 9A). Eguchi et al. found regulation of actin cytoskeleton to be the most significantly altered pathway in a proteomic analysis of plasma EVs isolated from four ME/CFS patients compared to patients with idiopathic chronic fatigue (ICF) or depression at baseline (Eguchi et al., 2020).

### Increased clusterin abundance post-exercise positively correlates with pain and fatigue

We found that the clusterin (CLU) 15min/0h ratio positively correlated with fatigue the morning of exercise (Figure 11B) and it was the only protein in which increased abundance 24 hours post-exercise positively correlated with both average severity of myalgia and arthralgia (Figure 9A). CLU was also one of the few proteins with decreased abundance 15 minutes post-exercise in controls (Figure 6A). CLU is a multifunctional molecular chaperone protein involved in a variety of physiological and pathologic processes, including clearance of cell debris, apoptosis, and assisting in the folding and conformational maturation of newly synthesized proteins as well as stress-denatured proteins (Poon et al., 2002).

Increased levels of clusterin were found in blood and brain tissues of patients with neurodegenerative diseases (Grewal et al., 1999; Ingram et al., 2014; Shepherd et al., 2020). CLU concentration in cerebrospinal fluid positively correlated with cognitive impairment severity, indicating a direct link with synaptic degeneration (Wang et al., 2020). Serum CLU has been associated with pain and inflammation and was increased in idiopathic inflammatory myopathy and early rheumatoid arthritis (Kropáčková et al., 2021; Kropáčková et al., 2020).

CLU levels in EVs were also elevated in myelodysplastic syndrome (Pecankova et al., 2022) and in multiple neurodegenerative diseases (Jiang et al., 2020). In our study, elevated CLU in EVs post-exercise is associated with worse myalgia, arthralgia, and fatigue indicating that this protein may have an important role in ME/CFS pathophysiology.

### Dysfunctional immune signaling in EVs in ME/CFS post-exercise

We found increased ANXA2, B2M and ORM1 in ME/CFS vs. controls 15 minutes post-exercise. These three proteins are members of the immune system protein pathways “antigen presentation”, “cytokine signaling”, and “adaptive immune system” (Figure 4B). Annexin II (ANXA2) is a calcium-dependent phospholipid-binding protein that orchestrates membrane repair, vesicle fusion, and cytoskeletal organization during the inflammatory response or tissue injury (Lim & Hajjar, 2021). ANXA2 is known to be involved in the biosynthesis of EVs and found in their cargo (Luo & Hajjar, 2013) and it has been shown that exosomal-ANXA2 promotes angiogenesis (Maji et al., 2017). ANXA2 in serum can increase inflammation by mediating macrophage M2 to M1 phenotypic change (Lin & Hu, 2017). Single-cell transcriptomics analysis has recently implicated enhanced macrophage differentiation from monocytes in ME/CFS pathophysiology (Ahmed et al., 2022).

Orosomucoid (ORM1) is an acute phase protein associated with fatigue. In our study, ORM1 had a significantly lower 24h/15min ratio in ME/CFS patients vs. controls, due to opposing trends in controls vs. patients (Figure 8, Supplementary Figure S3). A previous study found that the change in muscle soreness scores in healthy individuals positively correlated with serum ORM1 levels 24 hours post-exercise, suggesting that ORM1 is associated with the extent of exercise-induced damage and inflammation (Tékus et al., 2017). However, ORM1 is also upregulated in response to fatigue, leading to increased muscle glycogen levels and improved endurance. This positive feedback loop helps combat fatigue and maintain physiological homeostasis (Lei et al., 2016; Qin et al., 2016). Serum ORM levels correlated with fatigue severity in long-COVID patients, a disease with many overlapping symptoms with ME/CFS (Zavori et al., 2023).

Elevated serum total ORM (Sun et al., 2016) and ORM2 in cerebrospinal fluid have been found previously in ME/CFS patients vs. controls (Baraniuk et al., 2005). This upregulation of serum and CSF ORM at baseline may reflect the body’s attempt to mitigate the everyday fatigue that occurs in ME/CFS, while the altered dynamics of ORM1 in EVs post-exercise may indicate disruption of the muscle fatigue – ORM1 feedback loop.

### Members of the 14-3-3 protein family are increased in ME/CFS patients 15 minutes post-exercise

Increased DAPs 15 minutes post-exercise within the ME/CFS group were mainly associated with the apoptosis and cell cycle check points pathways, which include members of the 14-3-3 protein family (YWHAQ, YWHAG, YWHAB, YWHAZ) (Figure 6B). The Ingenuity pathway analysis showed that the 14-3-3 signaling pathway was significantly activated (z-score 2.5) in patients vs. controls at 15 minutes post-exercise (Figure 5). Three of these proteins (YWHAB, YWHAH, and YWHAQ) are expressed by the amygdala (Figure 7).

The 14-3-3 proteins are molecular chaperones widely expressed in all eukaryotic cells and especially in the central nervous system (CNS), accounting for about 1% of the total soluble brain protein (Berg et al., 2003). In the CNS, 14-3-3 proteins are usually induced in response to stresses, such as nerve damage and oxidation (Satoh et al., 2006). They play a key role in both development and neurodegeneration (Colucci et al., 2004; Morales et al., 2012; Shiga et al., 2006; Yacoubian et al., 2010). Interestingly, overexpression of YWHAQ enhanced the anti-apoptotic and anti-inflammatory effects of neural stem cell-derived EVs, which improved recovery after spinal cord injury (Rong et al., 2019). Furthermore, Eguchi et al. found YWHAH and YWHAQ were increased in abundance in the EVs of three ME/CFS patients compared to three controls (Eguchi et al., 2020). Higher levels of 14-3-3 protein family members in EVs in ME/CFS patients could be an attempt by the CNS to combat exertion-induced inflammatory signaling, which in our study was also increased in patients vs. controls.

### Limitations

The main limitation of this study is the small number of subjects, particularly for correlation analyses in which correlation coefficients are less stable with fewer samples. For that reason, we only presented correlations with |R| > 0.7. However, our study has more subjects than similar studies of EVs in a single group (Kobayashi et al., 2021; Whitham et al., 2018). Another key limitation is that we only included females. While the majority of ME/CFS patients are female, there are also male patients and it cannot be assumed that these results would apply to males as well, as many sex differences have been documented (Friedberg et al., 2023; Germain et al., 2022). A third limitation is that this study is limited to patients that were physically able to complete the exercise protocol, so the most severe and disabled ME/CFS patients are not represented in our study population. However, we did include mild and moderate ME/CFS patients with a wide range of symptom severity both on average and in response to exercise, which allowed us to find correlations between the EV proteome and symptom severity. The exercise test allowed us to synchronize the patients’ and sedentary controls’ molecular response to exertion, which enabled us to compare healthy controls’ and ME/CFS patients’ EV protein cargo dynamics post-exercise.

Regarding analysis of EVs, one limitation is that we did not isolate EVs from platelet-free plasma. While size exclusion chromatography removes platelets and larger microparticles from analysis, there is a possibility that platelets continued to release EVs *ex vivo* (Karimi et al., 2022) which can affect EV protein markers after exercise (McIlvenna et al., 2023). However, because we isolated plasma at three time points from the same individuals, we are still able to detect a specific effect of exercise. We also used the same methods for all samples and all subjects which allowed us to compare changes due to exercise across clinical groups.

## Conclusions

EV protein cargo changed differently in response to exercise in ME/CFS cases and sedentary controls. The changes in abundance post-exercise of several EV proteins in ME/CFS cases strongly correlate with the severity of certain symptoms. Dysregulated release of EVs and the presence of aberrant cargo relative to controls following exercise may play a role in the PEM experienced in females with ME/CFS.

## Acknowledgments

We are grateful to the individuals who participated in our strenuous exercise protocol. This research was supported by NIH NINDS/OD/NIDA/NHLBI/NHGRI through NINDS U54NS105541 to the Cornell Center for Enervating Neuroimmune disease and by a private donor. We thank Stephen Parry and Lynn Johnson at the Cornell Statistical Consulting Unit for their guidance on our statistical analysis. The REDCap database and blood processing at the Weill Cornell Medicine Clinical and Translational Science Center (CTSC) was supported by UL1 TR 002384 from the National Center for Advancing Translational Sciences of the NIH. We thank Carl Franconi, who supervised the biobank, REDCap survey database, and sample database. We thank the following individuals who participated in participant screening, and/or CPET, and/or blood collection: Victoria Birdsall, John Chia, Patricia Doty, Tiffany Ong, Betsy Keller, Susan Levine, Xiangling Mao, Geoffrey Moore, Maria Russell, Dikoma Shungu, Jared Stevens, Kristin Treat, and David Wang. We acknowledge the Cornell Proteomics and Metabolomics Facility, especially Ruchika Bhawal, Elizabeth T. Anderson, and Qin Fu, for technical assistance with the nano-LC MS/MS and preprocessing of the proteomics data.

