## Supplementary Figures S1-S3 for "Dysregulation of extracellular vesicle protein cargo in female ME/CFS cases and sedentary controls in response to maximal exercise"

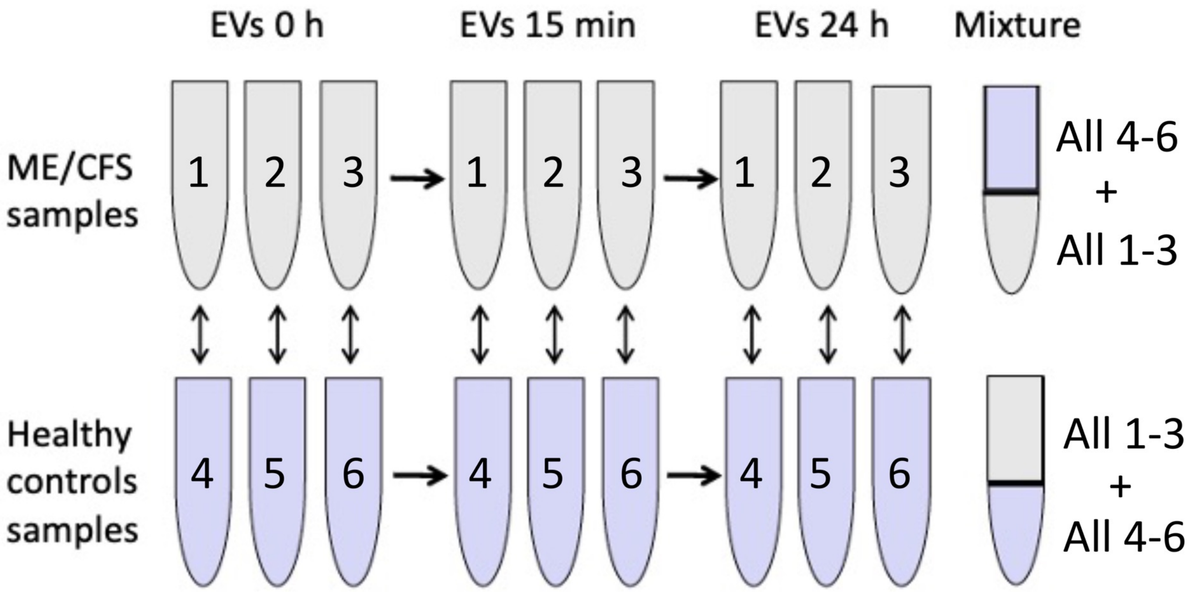


**Supplementary Figure S1: TMT10-plex analysis strategy.** 1, 2, and 3 represent three different ME/CFS subjects. 4, 5, and 6 represent three different healthy control samples.

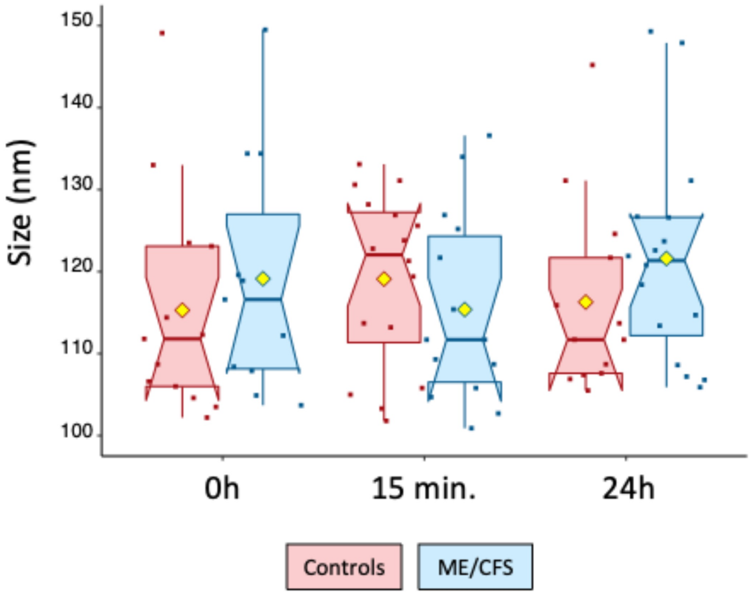


**Supplementary Figure S2: Nanoparticle Tracking Analysis.** Size of vesicles in nm in ME/CFS subjects and healthy controls. The yellow dot represents the mean. The non-parametric Wilcoxon signed-rank test was used to test the significance of differences ($p < 0.05$) between cases and controls. There were no significant differences.

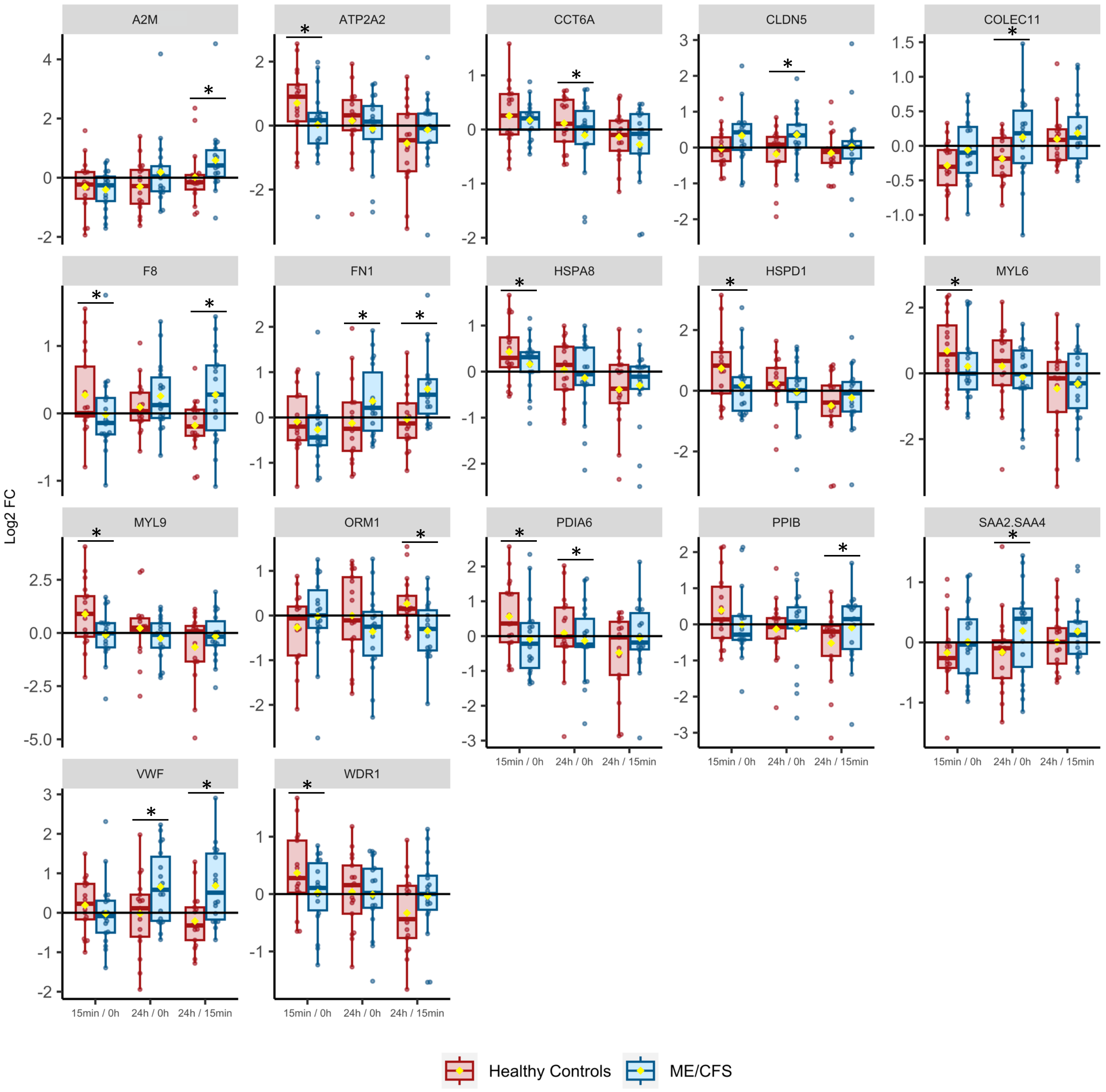


**Supplementary Figure S3: Within-subject fold changes over time for proteins changing differently post-exercise in ME/CFS patients vs. controls.** * shows significance between ME/CFS and control groups in the bootstrapping analysis (red dots in Figure 8), where a protein is significant if the FDR-adjusted confidence intervals for the median Log2FC of 10,000 bootstrapped datasets do not include 0 (q < 0.1). The yellow dot represents the mean.
