## Supplementary File S1 for "Dysregulation of extracellular vesicle protein cargo in female ME/CFS cases and sedentary controls in response to maximal exercise"

***MS method for Orbitrap Eclipse TMT-10plex analysis***

**Protein quantitation:**

Protein concentration for each sample was determined by gel quantification, each sample was run on a NOVEX 10% Bis/Tris mini-gel (Invitrogen, Carlsbad, CA) along with an *E. coli* lysates standard curve (0.5, 1, 2, 3 and 4 µg/lane). The SDS gel was visualized with SYPRORuby stain (Invitrogen), imaged by Typhoon 9400 scanner followed by ImageQuant TL 8.1 (GE Healthcare) software used for analysis.

**Protein digestion and TMT10-plex labeling**

Further processing of the proteins was then performed according to Thermo Scientific’s TMT Mass Tagging Kits and Reagents protocol (<http://www.piercenet.com/instructions/2162073.pdf> Note: TMT10plex Lot number: UG287008 was used for all sets) with a slight modification (refs#1-3). A total of 15 µg protein of each sample in the final buffer concentration: 50mM triethylammoniumbicarbonate (TEAB) pH 8.5, 6M Urea, 2M Thiourea, 1% SDS and 0.2% Triton X-100 was reduced with 10 mM tris(2-carboxyethyl)phosphine for 1 h at 34 °C, alkylated with 25 mM iodoacetamide for 45 min in the dark and then quenched with a final concentration of 60 mM Dithiothreitol (DTT). Each sample was digested separately using the S-Trap Micro Spin column (Protifi, Huntington NY) (ref# 4). After quenching, 12% phosphoric acid was added to a final concentration of 1% (v/v). Followed by 1:6 dilution (v/v) with 90% methanol, 0.1M TEAB pH 8.5. The sample were place into the S-Trap Micro spin column and centrifuged at 4000g for 30 sec. Then washed three times with 150 µl of 90% methanol, 0.1 M TEAB pH 8.5. Digestion was performed with 25 µl trypsin at 60 ng/µl (1:10 w/w) in 50 mM TEAB pH 8.5 added to the top of the spin column. The digest solution absorbed into the highly hydrophilic matrix, the spin columns were incubated overnight (16 hr) at 37 °C. Following incubation, the digested peptides were eluted off the S-Trap column sequentially with 40 µl each of 50 mM TEAB pH 8.5, followed by 0.2% formic acid, and finally 50% acetonitrile-0.2% formic acid. The eluates were pooled together and dried in speed vac. After drying down in the speed vac, each sample was reconstituted in 100 µl water and re-dried in speed vac to ensure that formic acid is removed.

Immediately before labeling, each sample was reconstituted into 30 µl 0.1M TEAB pH 8.5. The TMT 10-plex labels (0.2 mg dried powder) were reconstituted with 15 µl of anhydrous acetonitrile prior to labeling and added to each of the assigned 30 µl tryptic digest samples (1:13 w/w ratio peptide to TMT label) and incubated for 1 hour at room temperature. The labeled peptides from the 10 samples were pooled together as a set. The pooled peptides of each set were then evaporated to dryness and subjected to cleanup by solid phase extraction (SPE) on MCX Cartridges (Waters, Milford, MA). Labeling incorporation was then verified using Orbitrap Eclipse Thermo-Fisher Scientific, San Jose, CA). The eluted tryptic peptides were evaporated to dryness, and ready for the fractionation via a high pH reverse phase peptide fractionation kit as described below.

**High pH Reversed-Phase Peptide Fractionation**

Fractionation was carried out using a Pierce™ High pH Reversed-Phase Peptide Fractionation Kit (Thermo-Fisher Scientific, San Jose, CA) following provided vendor protocol. Briefly, after the protective white tip was removed and the spin column was centrifuged at 5000 g for 2 min to remove storage solution, it was then conditioned with 300 µl ACN followed by two washes with 300 µl 0.1% TFA. 80 µg tryptic TMT labeled peptides in 0.1% TFA was loaded onto the spin column and then washed with 300 µl water. The TMT labeled peptides were step eluted with 10%, 12.5%, 15%, 17.5%, 20%, 22.5%, 25%, 50% and 80% ACN in 0.1% triethylamine respectively; The nine eluted fractions were then pooled into three fractions and dried down in the speed vac. Finally, each fraction was reconstituted into 66 µl of 2% ACN/0.5% FA for subsequent nanoLC-MS/MS analysis.

**Nano-scale reverse phase chromatography and tandem MS (nanoLC-MS/MS)**

The nanoLC-MS/MS analysis was carried out using an Orbitrap Eclipse (Thermo-Fisher Scientific, San Jose, CA) mass spectrometer equipped with a nanospray Flex Ion Source using real-time search SPS MS3 method coupled with the UltiMate3000 RSLCnano (Dionex, Sunnyvale, CA). Each reconstituted fraction (5 μL for global proteomics fractions) injected onto a Thermo Acclaim PepMap 100 nanoViper trap column (3um, 100um x 20mm, Dionex) at 20 μL/min flow rate for on-line desalting, and separated on a Thermo Acclaim PepMap RSLC nanoViper column (2um, 75um x 25cm). The column was equilibrated with 2% acetonitrile (ACN) in 0.1% aqueous formic acid (eluant A) prior to each run. The labeled peptides were eluted in a 120 min gradient of 5% to 33% eluant B containing 95% ACN in 0.1% formic acid at 300 nL/min, followed by an 8-min ramping to 90% B, a 9-min hold and 25min re-equilibration with 2% ACN-0.1% FA prior to the next run. The Orbitrap Eclipse is operated in positive ion mode with nano spray voltage set at 1.9 kV and source temperature at 300 °C. External calibration for FT, IT and quadrupole mass analyzers was performed. For global proteomics fractions, the instrument was operated using a RTS SPS MS3 workflow based on the manufacturer’s provided template. Specifically, the RTS MS3 template contained 2.5 second “Top Speed” data-dependent CID-MS/MS scans (for peptide identifications by RTS) followed by SPS of 10 MS^2^ product ions for subsequent MS^3^ in FT. In RTS mode, the HomoSapiens NCBI downloaded on 3/14/2019 that has about 81268 entries was imported as the FASTA database with trypsin as the enzyme for real-time spectral database search. The search parameters included: TMT 6-plex modifications on lysine and N-terminal amines (∆mass 229.1629), carbamidomethyl modification of cysteine (∆mass 57.0215), and maximum 2 variable methionine oxidation per peptide, and 1 missed cleavage. A maximum search time for 35 ms allowed for the RTS MS^3^ searching. The MS^3^ scan was carried out using a mass range of 100-500 m/z, an MS isolation window of 1.1 m/z and MS^2^ isolation window of 2.0 m/z were used. A resolving power of 50,000 with a normalized collision energy of 55% for peptide quantitation. Other parameters included 500% normalized AGC target and 118 ms for maximum injection time. Dynamic exclusion parameters were set at 1 within 50s exclusion duration with ±10 ppm exclusion mass window. All data were acquired under Xcalibur 4.3 operation software in Orbitrap Eclipse (Thermo Fisher Scientific).

**Data processing, protein identification and data analysis**

All raw MS spectra were processed and searched using the Sequest HT search engine within the Proteome Discoverer 2.3 (PD2.3, Thermo). The MS/MS spectra were searched against HomoSapiens NCBI downloaded on 3/14/2019 that has about 81268 entries. Oxidation of M, deamidation of N and Q, and acetylation of protein N-terminus were specified as dynamic modifications of amino acid residues; carbamidomethyl C and TMT 6 plex on N-terminal and Lys residues were specified as a static modification. The default search settings used for 10-plex TMT quantitative processing and protein identification in PD2.3 searching software were: two mis-cleavage for full trypsin with fixed carbamidomethyl modification of cysteine, fixed 6-plex TMT modifications on lysine and N-terminal amines and variable modifications of methionine oxidation, deamidation on asparagines/glutamine residues and protein N-terminal acetylation. The peptide mass tolerance and fragment mass tolerance values were 10 ppm, 0.6 Da for MS2 and 20 ppm for MS3, respectively. Identified peptides were further filtered for maximum 1% FDR using the Percolator algorithm in PD 2.3 along with additional peptide confidence set to high and peptide mass accuracy ≤5 ppm. The TMT10-plex quantification method within Proteome Discoverer 2.3 software was used to calculate the reporter ion abundances in MS3 spectra that were corrected for the isotopic impurities. Both unique and razor peptides were used for quantitation. Signal-to-noise (S/N) values were used to represent the reporter ion abundance with a co-isolation threshold of 50% and an average reporter S/N threshold of 10 and above required for quantitation spectra to be used. The S/N values of peptides, which were summed from the S/N values of the PSMs, were summed to represent the abundance of the proteins. For relative ratio between the two groups, normalization on total peptide amount for each sample was applied. The search result including ratio, p value, peptide abundance for each sample was output to Microsoft Excel software for further data analysis.

**Acknowledgements**

We thank the Proteomic and Metabolomics Facility of Cornell University for generating the mass spectrometry data and the funding support from HHMI Transformative Technology 2019 program for the Orbitrap Eclipse system.
